# Hypoglycosylation lowers the mechanical activation threshold of Piezo1 and enhances cortical neuronal mechanotransduction: implications for PMM2-CDG

**DOI:** 10.64898/2026.08.03.742511

**Authors:** Albert Edo-Pérez, Gorane Rodríguez-Urquirizar, Alicia Fernández-Arroyo, Julia Carrillo-García, José M. Fernández-Fernández

## Abstract

Piezo1 is a mechanically activated cation channel whose N-linked glycans support protein maturation and plasma membrane trafficking, but their contribution to mechanical gating is unknown. We asked whether hypoglycosylation alters Piezo1 mechanosensitivity and cortical neuronal mechanotransduction, with potential relevance to neurological manifestations of congenital disorders of glycosylation (CDG). Human Piezo1 was studied in HEK293 cells after mutation of two conserved cap-domain N-glycosylation sites or inhibition of N-glycan maturation with swainsonine or kifunensine. Mechanically activated currents were recorded by cell-attached patch-clamp during incremental negative-pressure pulses, whereas Ca^2+^ responses were measured during uniaxial stretch. Piezo1 abundance, synaptic localisation and stretch-evoked Ca^2+^ signals were also examined in primary mouse cortical neurons. On poly-L-lysine, N2293Q or N2330Q shifted the pressure-response relationship towards lower activating pressures without changing maximal current or inactivation kinetics. This effect was absent on collagen. Swainsonine and kifunensine reduced mature Piezo1 glycosylation and lowered the mechanical activation threshold. Hypoglycosylation enhanced Ca^2+^ entry during submaximal stretch in HEK293 cells. In cortical neurons, inhibition of glycan maturation increased somatic Piezo1 immunoreactivity without changing its association with synaptic markers, and potentiated Ca^2+^ responses to both the Piezo1 activator Yoda1 and submaximal stretch. Thus, mature N-glycans and the extracellular adhesive environment jointly set Piezo1’s mechanical activation threshold rather than merely regulating biosynthesis and trafficking. These findings establish glycosylation-mechanics coupling as a determinant of neuronal force sensing and suggest that, by facilitating Piezo1 recruitment, defective glycosylation may increase cortical vulnerability to mechanical stress, potentially contributing to head trauma-triggered neurological episodes in phosphomannomutase 2 deficiency (PMM2-CDG).

**Key points:**

- Piezo1 channels convert mechanical forces into electrical and calcium signals. N-linked glycans support channel trafficking to the plasma membrane, but whether they tune the force needed for Piezo1 activation was unknown.
- Mutating either of two conserved N-glycosylation sites in Piezo1 cap domain, or pharmacologically restricting N-glycan maturation, lowered channel’s mechanical activation threshold without changing maximal current or inactivation.
- This sensitisation depended on the adhesive substrate (occurred on poly-L-lysine but not collagen), and was most evident during submaximal stretch, showing that glycosylation and the extracellular mechanical environment jointly determine Piezo1 force sensing.
- In mouse cortical neurons, impaired N-glycan maturation increased somatic Piezo1 abundance and enhanced Ca^2+^ responses to its chemical activator Yoda1 and stretch, without changing synaptic localisation.
- By allowing weak mechanical inputs to recruit Piezo1 more effectively, defective glycosylation may increase cortical responses to mechanical stress and help explain susceptibility to head trauma-triggered neurological episodes in phosphomannomutase 2 deficiency (PMM2-CDG).

## Introduction

Glycosylation is one of the most important post-translational modifications that takes place in the endoplasmic reticulum (ER) and Golgi apparatus. It is essential for the proper folding of transmembrane proteins, enhancing their stability and regulating their function (Stanley, 2024). N-glycosylation, a major glycosylation pathway, involves the attachment of glycans to asparagine residues in proteins, within the consensus sequence Asn-X-Ser/Thr (where X is any amino acid except proline) (Marshall & Neuberger, 1964; Marshall & Neuberger,1970). This process occurs in two phases: 1) the oligosaccharide synthesis on a lipid carrier at the ER membrane; and 2) the subsequent processing of N-glycans by glycosidases and glycosyltransferases within the ER lumen and Golgi apparatus (Reily et al., 2019). The resulting N-glycans can be classified as high mannose, hybrid or complex N-glycans, with the latter two representing the more mature forms (Stanley et al., 2022; Stanley, 2024).

Phosphomannomutase 2 (PMM2) plays an indispensable role in the early stages of the N-glycosylation (Sharma et al, 2014). Loss-of-function mutations in the *PMM2* gene result in Phosphomannomutase 2 Deficiency (PMM2-CDG), the most prevalent congenital disorder of N-linked glycosylation (Jaeken et al., 2009; Francisco et al., 2023). PMM2-CDG affects multiple organ systems, with severe neurological alterations significantly impairing patient’s quality of life. The cerebellum is consistently affected in PMM2-CDG patients, showing progressive atrophy (Freeze et al., 2012; de Diego et al., 2017). In addition, stroke-like episodes (SLEs) are a common, unpredictable, and severe neurological complication observed in PMM2-CDG patients (Freeze et al., 2012; Izquierdo-Serra et al., 2018). SLEs also complicate paroxysmal neurological diseases such as familial hemiplegic migraine (FHM), predominantly caused by mutations in the *CACNA1A* gene, which encodes the pore-forming α_1A_ subunit of the neuronal Ca_V_2.1 channel (Pietrobon, 2013).

The mechanisms underlying both cerebellar syndrome and SLEs in PMM2-CDG remain poorly understood. Notably, a striking similarity has been reported between the clinical, neuroimaging and neurophysiological features of PMM2-CDG patients and those with *CACNA1A* mutations, including not only SLEs but also ataxia, eye movement abnormalities, and cerebellar atrophy (Izquierdo-Serra et al., 2018). Furthermore, hypoglycosylation of Ca_V_2.1 channel subunits has been shown to induce gain-of-function effects on channel gating, which mirror those reported for pathogenic *CACNA1A* mutations linked to FHM and some forms of ataxia with cerebellar atrophy (Izquierdo-Serra et al., 2018). This suggests a potential link between hypoglycosylation of Ca_V_2.1 channels and neurological manifestations of PMM2-CDG, offering insight into possible prophylactic or therapeutic strategies for these acute and stressful neurological complications (Martínez-Monseny et al., 2019).

Mild cranial trauma has been identified as a potential trigger for SLEs in PMM2-CDG patients (Izquierdo-Serra et al., 2018; Wicker et al., 2023). The brain, being a mechanosensitive organ, can transduce mechanical forces into a variety of neurological responses (Gaetz, 2004; Tufail et al., 2010; Tyler, 2012). Mechanosensitive ion channels have been proposed as key players in this transduction process. Specifically, mechanosensitive cationic currents activated in the soma are coupled to the triggering of action potentials in central neurons of the mammalian brain, including those in the cortex and hippocampus (Nikolaev et al., 2015). Although the molecular nature of these mechanosensitive channels remains unclear, members of the Piezo family channel are promising candidates. Piezo channels are known to mediate mechanically activated cation currents and are expressed in the mammalian brain (Coste et al., 2010). Additionally, brain mechanosensitive channels exhibit properties such as single channel conductance, stretch sensitivity, and low cation selectivity, consistent with the characteristics of Piezo channels (Coste et al., 2010; Nikolaev et al., 2015). Moreover, Piezo1 has been identified as an N-glycosylated protein on the surface of mouse embryonic stem cells and neural progenitors (Wollscheid et al., 2009), with its glycosylation status shown to influence its functional expression (Li, Ng et al., 2021).

In this study, we investigate whether hypoglycosylation of Piezo1 alters its channel activity in a way that could contribute to the neurological symptoms observed in PMM2-CDG, particularly the occurrence of SLEs triggered by mechanical brain stimulation following head trauma. Using electrophysiological recordings, Ca^2+^ imaging, mutagenesis of Piezo1 N-glycosylation sites, and mature N-glycosylation inhibitors such as swainsonine and kifunensine, we demonstrate that hypoglycosylation of Piezo1 heterologously expressed in HEK293 cells increases its mechanosensitivity in response to both negative pressure and cellular stretch, without affecting channel inactivation kinetics. This enhancement of mechanosensitivity is substrate-dependent, occurring in the presence of poly-L-lysine but not collagen. Additionally, inhibition of mature N-glycosylation in murine cortical neurons increases Piezo1 levels at the soma without altering its synaptic localisation. The N-glycosylation inhibitors also enhance neuronal Ca^2+^ responses upon Piezo1 stimulation, both via its chemical activator Yoda1 and mechanical stretch. Taken together, our findings suggest that hypoglycosylation-induced gain-of-function in Piezo1 may contribute to heightened brain sensitivity to mechanical stimuli, potentially exacerbating the neurological manifestations of PMM2-CDG.

## Methods

### cDNA constructs

The cDNA of the wild-type (WT) human Piezo1 (hPiezo1) channel (isoform containing the polymorphic deletion of residue K1878) was originally cloned into a pIRES2-EGFP vector and generously provided by Dr. Frederick Sachs (Department of Physiology and Biophysics, State University of New York at Buffalo, Buffalo, NY, United States of America). Piezo1 N2293Q and N2330Q mutants (equivalent to those previously described as N2294Q and N2331Q by Li, Ng et al, 2021) were generated through site-directed mutagenesis of the WT hPiezo1 cDNA (GenScript Corporation, Piscatway, NJ, USA). All cDNA clones were fully sequenced to ensure integrity.

### Heterologous expression in HEK293 cells

HEK293 cells were maintained in Dulbecco’s modified Eagle’s medium (L0102, Biowest) supplemented with 10% foetal bovine serum (S181H, Biowest) and 100 U/mL Penicillin/Streptomycin (15140-122, Gibco). HEK293 cells were seeded on 6 well plates at 70-80% confluency and transfected with 3 μg of DNA using JetPEI (101-10N, Polyplus) or Lipofectamine 3000 (L3000001, Invitrogen). The same day of transfection, treatments with 250 μM swainsonine (3208, Tocris) or 25 μM kifunensine (3207, Tocris) were applied and maintained for 48 hours, when indicated.

At 24 hours post-transfection, HEK293 cells were washed twice with PBS and mechanically detached and collected in PBS. Cells were centrifuged at 335 g for 5 minutes and seeded on 35-mm petri dishes (10035, SPL Life Sciences) previously coated with 5 μg/cm^2^ collagen (354236, Corning) or 6 μg/cm^2^ Poly-L-Lysine (P6282, Sigma-Aldrich) for 1h at 37°C, for electrophysiological recordings during the following day. In the case of mechanical stretching experiments, cells were seeded at 500.000 cells per stretching chamber, which were previously coated with 18.75 μg/cm^2^ Poly-L-Lysine (P6282, Sigma-Aldrich) overnight at 37°C.

### Ethical approval and animal experiments

All animal experiments were conducted in accordance with the Spanish legislation on animal protection, approved by the local animal care committee (Comité Ético de Experimentación Animal del Consorci Parc de Recerca Biomèdica de Barcelona (CEEA-PRBB)), including Universitat Pompeu Fabra; Approved Procedure: PML16-0021PR1-P1), and conformed to the Directive 2010/63/EU of the European Parliament on the protection of animals used for scientific purposes.

### Isolation and primary culture of murine cortical neurons

Cortical neurons were isolated from E18 mouse embryos. Pregnant mice were euthanised by CO_2_ inhalation in accordance with the directives of the Council of the European Communities N. 86/609/CEE. Embryos were rendered hypothermic and decapitated. Brains were extracted and cortices were isolated under a Leica S6 E stereomicroscope in ice-cold extraction medium (HBSS 1X, 4.5% Glucose, 1% Penicillin/Streptomycin) (Gibco and Sigma). Meninges were carefully removed, and hippocampi were detached from the cortical tissue.

Isolated cortices were chopped into 4-5 pieces and enzymatically dissociated with 0.05% trypsin (P6407, Sigma) for 17 min at 37°C, 5% CO_2_. The tissue was then mechanically dissociated by gentle pipetting and filtered through a 70 μm pore diameter cell strainer (141379C, Cultek). Cells were seeded on poly-L-lysine (50 μg/mL; 25300-054, Gibco) coated coverslips (174950, Thermo Fisher Scientific) (125,000 cells/coverslip) or stretch chambers (SC-0040, Strex) (300,000 cells/chamber), in plating medium (DMEM High Glucose 1X, 10% Horse Serum, 1% Penicillin/Streptomycin) (Gibco), which was replaced with Neurobasal Complete (NBC) medium (Neurobasal medium 1X, B27 supplement 1X, GlutaMAX 1X, 1% Penicillin/Streptomycin; Gibco) after 2 hours.

To inhibit glial cell proliferation, cultures were treated with 2 μM cytosine β-D-arabinofuranoside (AraC) (C1768, Sigma) on day in vitro 3 (DIV3). Neurons were maintained in culture for 10-15 days before use in experiments.

When inhibition of mature glycosylation was required, neurons were treated with 250 μM swainsonine (3208, Tocris) or 25 μM kifunensine (3207, Tocris) for 72h before experimental recordings. The working concentration for each inhibitor was determined based on previous reports using swainsonine (Fiaux et al., 2006; Cholich et al., 2020) and kifunensine (Elbein et al., 1991; Walker et al., 2010) on human cell lines. Kifunensine inhibits ER and Golgi α-mannosidases, which prevent the trimming of the α2Man residues in the Man_8_GlcNAc_2_ high-mannose N-glycan, therefore blocking the synthesis of both hybrid and complex N-glycans (Elbein et al., 1990). Swainsonine inhibits the Golgi α-mannosidase II and the subsequent trimming of the terminal α3Man and α6Man residues in the Man_5_GlcNAc_2_ glycan, which prevents the formation of complex N-glycans (Tulsiani & Touster, 1983).

### Western blotting

After 48 hours of treatment with either vehicle (DMSO, 1:1000), 250 μM swainsonine, or 25 μM kifunensine, HEK293 cells transfected with the WT hPiezo1 cDNA were lysated on ice in using 40μL of lysis buffer (10mM Tris-Base, 1 mM EDTA, 140mM NaCl, 0.1% Sodium deoxycholate, 0.1% SDS, 1% Triton-X-100) supplemented with 10% protease inhibitor cocktail (11836153001, Roche), 1mM NEM, 1mM PMSF and 2mM TCEP. Lysates were incubated during 10 min at 4°C on a rotating wheel and centrifuged at 4°C for 20 min at 13.000 g. Protein concentration of cleared lysates were determined using Pierce BCA protein assay kit (Thermo Fisher Scientific).

When indicated, samples (15 μg) were digested with PNGase (P0704S, NEB) for 1h on ice following manufacturer’s protocol. Samples were mixed with NuPage LDS sample buffer (NP0007, Thermo Fisher Scientific) and NuPage sample reducing agent (NP0004, Thermo Fisher Scientific) and run on a 3-8% Tris-Acetate gel (EA0375PK2, Thermo Fisher Scientific). Proteins were then transferred to a nitrocellulose membrane (1794158, Bio-Rad) and incubated with blocking solution containing 5% Bovine serum albumin in TTBS for 1h RT. Membranes were then incubated with anti-Piezo1 (1:500, NBP2-75617, Novus Biologicals) and anti-alpha-actinin (1:2000, sc-166524, Santa Cruz) diluted in blocking solution overnight at 4°C in a tube roller. Secondary antibody anti-mouse (1:2000, M2650, Sigma) was diluted in blocking solution and incubated for 1h RT.

Detection was performed using enhanced chemiluminescence detection kit (34580, Thermo Fisher Scientific).

### Electrophysiological recordings

Cell-attached recordings were performed 48 hours post-transfection using borosilicate glass patch pipettes (1.8 to 2.2 MΩ) filled with a solution containing 130 mM NaCl, 5 mM KCl, 1 mM CaCl_2_, 1 mM MgCl_2_, 10 mM tetraethylammonium-Cl and 10 mM HEPES (adjusted to pH 7.3 with Tris-base and to 300 mOsm with mannitol). Bath solution contained 140 mM KCl, 1mM MgCl_2_, 10 mM glucose and 10 mM HEPES (adjusted to pH 7.3 with Tris-base and to 310 mOsm with mannitol). Piezo1 channel stimulation was performed using a High-Speed Pressure Clamp (HSPC-1, ALA Scientific Instruments) with increasing pulses of negative pressure from −10 to −80 mmHg (ΔP = −10 mmHg). Voltage was clamped at −80 mV using an EPC10-USB patch-clamp amplifier (HEKA Elektronik). Data were analysed using MATLAB (MathWorks) and Igor Pro 8 (WaveMetrics).

The half-maximal activating pressure (AP_50_) was obtained using Igor Pro 8 (WaveMetrics) by fitting a double exponential into the I/I_max_ data and interpolating the pressure value where I/I_max_ = 0.5.

The time constant of inactivation (τ_inact_) was obtained using MATLAB by fitting the traces to a double exponential function between the peak current and the end of the stimulation according to the following double exponential equation:

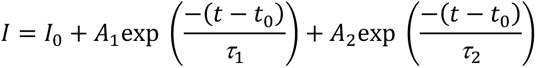

The double exponential function provides a better fit to the biphasic inactivation kinetics of Piezo1, consisting of a fast initial component followed by a much slower second component. The inactivation kinetics of Piezo1 will be reported as the time constant of inactivation of the fast component (τ_1_).

### Calcium imaging

At 48 hours post-transfection, HEK293 cells were washed twice with imaging solution containing 140 mM NaCl, 0.5 mM MgCl_2_, 1.2 mM CaCl_2_, 2.5 mM KCl, 10 mM HEPES and 5 mM Glucose (adjusted to pH 7.3 with Tris-base and to 305 mOsm with mannitol) and incubated with 400 μL of 10 μg/mL Calbryte630 AM (20720, AAT Bioquest) for 25 minutes at 37°C, 5% CO_2_. HEK293 cells were then washed three times with imaging solution and mounted to the microscope. A similar protocol was used for primary cortical neurons from mice, with slight differences: experiments were performed at DIV10-15, the imaging solution contained 140 mM NaCl, 1 mM MgCl_2_, 2.5 mM CaCl_2_, 3 mM KCl, 10 mM HEPES and 10 mM Glucose (adjusted to pH 7.3 with Tris-base and to 305 mOsm with mannitol), and cells were incubated with 400 μL of 10 μg/mL Calbryte520 AM (20650, AAT Bioquest) for 20 minutes at 37°C, 5% CO_2_.

To apply mechanical stimuli, stretching chambers containing the cells were mounted into a uniaxial cell stretching system (STB-150, Strex) and two 500 ms pulses of 40% and 80% stretching (relative to the initial chamber length) were applied with a delay of 10 minutes. Fluorescence data (F575 for Calbryte630 AM and F480 for Calbryte520 AM) were acquired with either HCImage or Aquacosmos imaging processing software (Hamamatsu Photonics). Data were normalised to the signal measured before stretch stimulation.

For Ca^2+^ imaging recordings in response to chemical stimulation of Piezo1 channels with Yoda1, coverslips containing the murine primary cortical neurons at DIV10-15 were washed with the imaging solution and incubated with Calbryte520 AM (20650, AAT Bioquest), as described for neuronal stretching experiments. The coverslips were then mounted to the microscope with a R-25 bath recording chamber (Warner Instruments). After a 2-minute baseline, neurons were perfused with 10 μM Yoda1 (5586, Tocris) for 30 seconds. During the first 3 minutes of recording, neurons were exposed to 1 μM TTX (BML-NA120-0001, Enzo LifeSciences). Fluorescence data (F480) were acquired with HCImage imaging processing software (Hamamatsu Photonics) and were normalised to the signal measured before Yoda1 stimulation.

### Confocal microscopy

For Piezo1 localisation analysis, murine cortical neurons were fixed with 4% PFA supplemented with 1% sucrose for 10 min at RT, permeabilised with 0.2% Triton X-100 (Sigma) in PBS for 10 min at RT, and blocked in 5% BSA in PBS for 45 minutes at RT. Cells were then sequentially incubated with primary antibodies (Table 1) overnight at 4°C in a humid chamber, secondary antibodies (Table 1) for 45 minutes at RT, and DAPI (1:2.000 in PBS; 62248, Thermo Fisher Scientific) for 10 min at RT, all diluted in 1% BSA in PBS. Cells were imaged with a Leica TCS-SP8 confocal microscope with a 63X 1.40 immersion oil objective. Image processing was performed with ImageJ software (National Institute of Health). Piezo1, vGlut1, PSD95, vGAT and Gephyrin puncta masks and Manders Overlap Coefficient were generated using Cell Profiler (Stirling et al., 2021). Mean intensity values were quantified using ImageJ (National Institute of Health).

**Table 1.** List of antibodies used in immunostainings of primary cortical neurons from mice.

|  | Reference | Supplier | Dilution |
| --- | --- | --- | --- |
| **Primary antibodies** |  |  |  |
| Gephyrin | 147 011 | Synaptic Systems | 1:500 |
| Piezo1 | NBP1-78446 | Novus Biologicals | 1:350 |
| PSD95 | 124 308 | Synaptic Systems | 1:500 |
| vGAT | 131 004 | Synaptic Systems | 1:500 |
| vGlut1 | 135 011 | Synaptic Systems | 1:500 |
| **Secondary antibodies** |  |  |  |
| Alexa Fluor 488 Goat Anti-Guinea Pig IgG (H+L) | A-11073 | Life Technologies | 1:2.000 |
| Alexa Fluor 647 Donkey Anti-Mouse IgG (H+L) | A-31571 | Life Technologies | 1:2.000 |
| Alexa Fluor 555 Goat Anti-Rabbit IgG (H+L) | A-21429 | Life Technologies | 1:2.000 |

### Statistical analysis

Statistical analysis and plots were performed using GraphPad Prism. Normality was assessed with the Shapiro-Wilk test. The Mann-Whitney U test, one-way ANOVA followed by Dunnett’s post-hoc test, or Kruskal-Wallis followed by Dunn’s post-hoc test were applied as appropriate. Data are shown as mean ± SEM. Statistically significant differences were considered as follows: *P < 0.05, **P < 0.01, ***P < 0.001 and ****P < 0.0001.

## Results

### Substrate-Dependent Modulation of Piezo1 Mechanosensitivity by N-Glycosylation

Li, Ng et al. (2021) demonstrated that two conserved asparagine residues (N2294 and N2331) in the cap domain of Piezo1 are critical for full N-glycosylation and proper channel trafficking. Their findings confirmed that the double mutant N2294Q/N2331Q exhibited minimal plasma membrane localisation and negligible stretch-activated currents, in stark contrast to the robust membrane localisation and mechanosensitive activity of the wild-type (WT) channel. However, partial glycosylation at either N2294 or N2331 proved sufficient to maintain functional trafficking, enabling systematic evaluation of N-glycosylation’s role in Piezo1 activity (Li, Ng et al., 2021). Notably, while Li, Ng et al. (2021) reported no impact of single glycosylation-site mutations on channel inactivation kinetics, the functional consequences of hypoglycosylation on Piezo1 mechanoactivation remained unexplored. To address this gap, we compared the mechanical activation profiles of single glycosylation mutants (N2294Q and N2331Q) with WT Piezo1 channels heterologously expressed in HEK293 cells. This study employed a human Piezo1 (hPiezo1) isoform harbouring the K1878 polymorphic deletion, which shifts residue numbering. Consequently, the corresponding N-glycosylation site mutants analysed were N2293Q and N2330Q.

It is well established that substrate stiffness and composition significantly influence the mechanical properties of Piezo1 (McHugh et al., 2012; Qi et al., 2015; Marchioni et al., 2021; Raha et al., 2023). Therefore, to evaluate the impact of hypoglycosylation on Piezo1 mechanosensitivity, we tested two common surface coatings: poly-L-lysine (PLL) and collagen.

Cell-attached patch-clamp recordings were performed on HEK293 cells expressing WT hPiezo1, with incremental negative pressure pulses applied via the recording pipette. For WT channels, peak current amplitudes (I_max_) showed no significant difference between PLL-coated (−157.0 ± 23.08 pA, n = 20) and collagen-coated substrates (−116.4 ± 10.87 pA, n = 34; P = 0.132) (Fig. 1*A* and *B*). However, substrate type markedly affected mechanosensitivity, as evidenced by the half-maximal activating pressure (AP_50_). On PLL, AP_50_ was −20.53 ± 2.26 mmHg (n = 20), whereas collagen reduced it to −15.12 ± 1.18 mmHg (n = 34; P = 0.044), indicating enhanced sensitivity on collagen (Fig. 1*C* and *D*).

**Figure 1.**
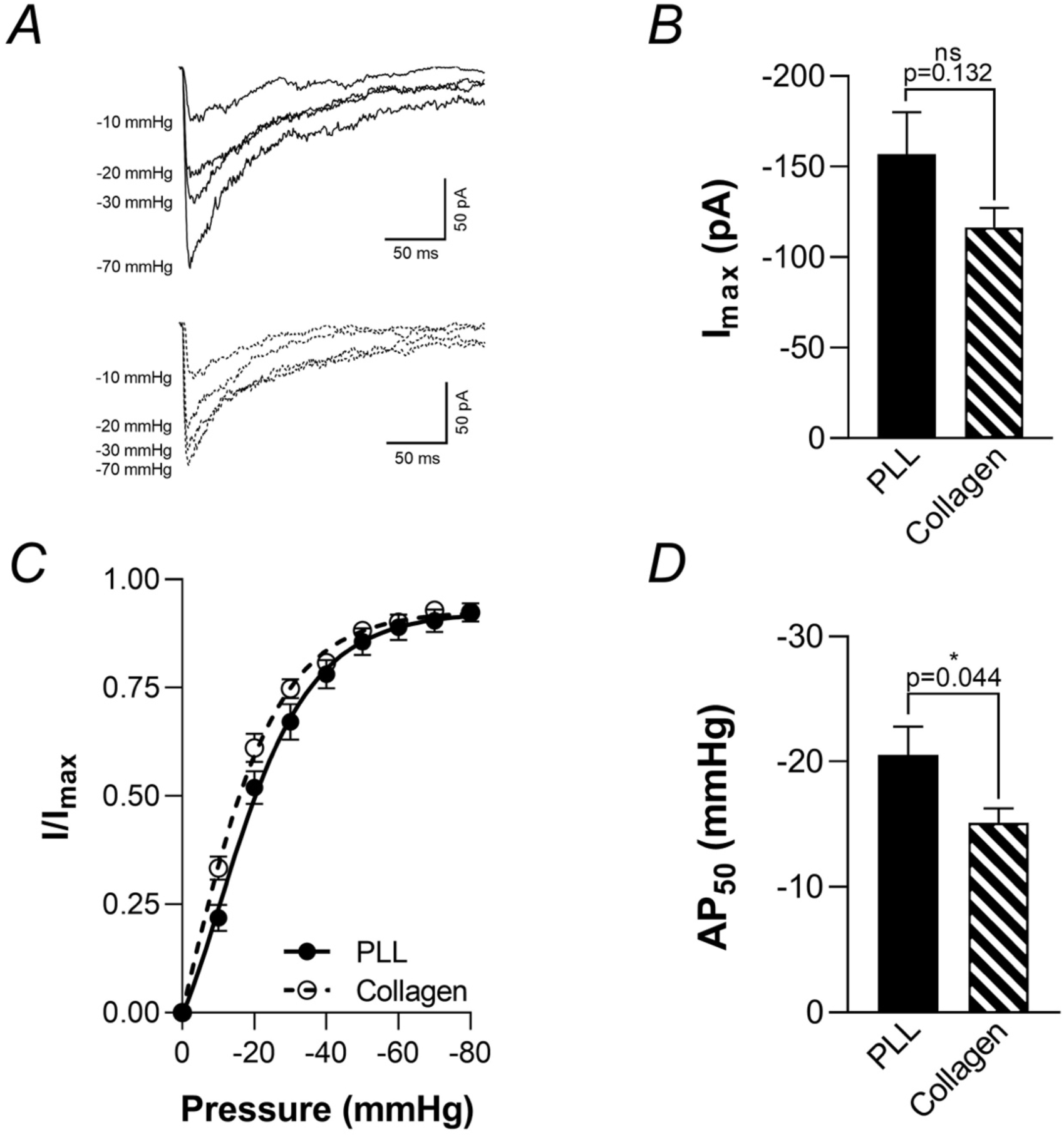
Mechanosensitivity of the hPiezo1 WT channel depends on the surface coating agent. *A*, representative traces of mechanically activated currents obtained from cell-attached patches on HEK293 cells heterologously expressing the human Piezo1 (hPiezo1) WT channel on PLL- (top) or collagen-coated (bottom) substrates. Only traces in response to −10, −20, −30 and −70 mmHg pulses are shown for visualisation purposes. *B*, quantification of the peak current per patch (I_max_) elicited by negative pressure pulses. *C*, normalised current-pressure relationship calculated as the I/I_max_ ratio. *D*, average half-maximal activating pressure (AP_50_) for each experimental condition. Data presented as mean ± SEM. Statistical significance was assessed using Mann-Whitney’s U test. (*P < 0.05; ns, not significant; exact P values shown; sample sizes: PLL, n = 20; Collagen, n = 34).

In parallel we also run the same experiments using HEK293 cells heterologously expressing either N2293Q or N2330Q hPiezo1 mutants. In collagen-coated substrates, neither N2293Q (−115.9 ± 9.67 pA, n = 25) nor N2330Q (−119.5 ± 21.82 pA, n = 13) mutants differed significantly from WT (-116.4 ± 10.87 pA, n = 34, P >0.999) in I_max_ (Fig. 2*A* and *B*). Similarly, AP_50_ values for N2293Q (−15.45 ± 1.61 mmHg, n = 25) and N2330Q (−13.93 ± 2.55 mmHg, n = 13) aligned with WT (−15.12 ± 1.18 mmHg, n = 34) (P = 0.985 and P = 0.894, respectively), suggesting preserved mechanosensitivity despite hypoglycosylation (Fig. 2*C* and *D*).

**Figure 2.**
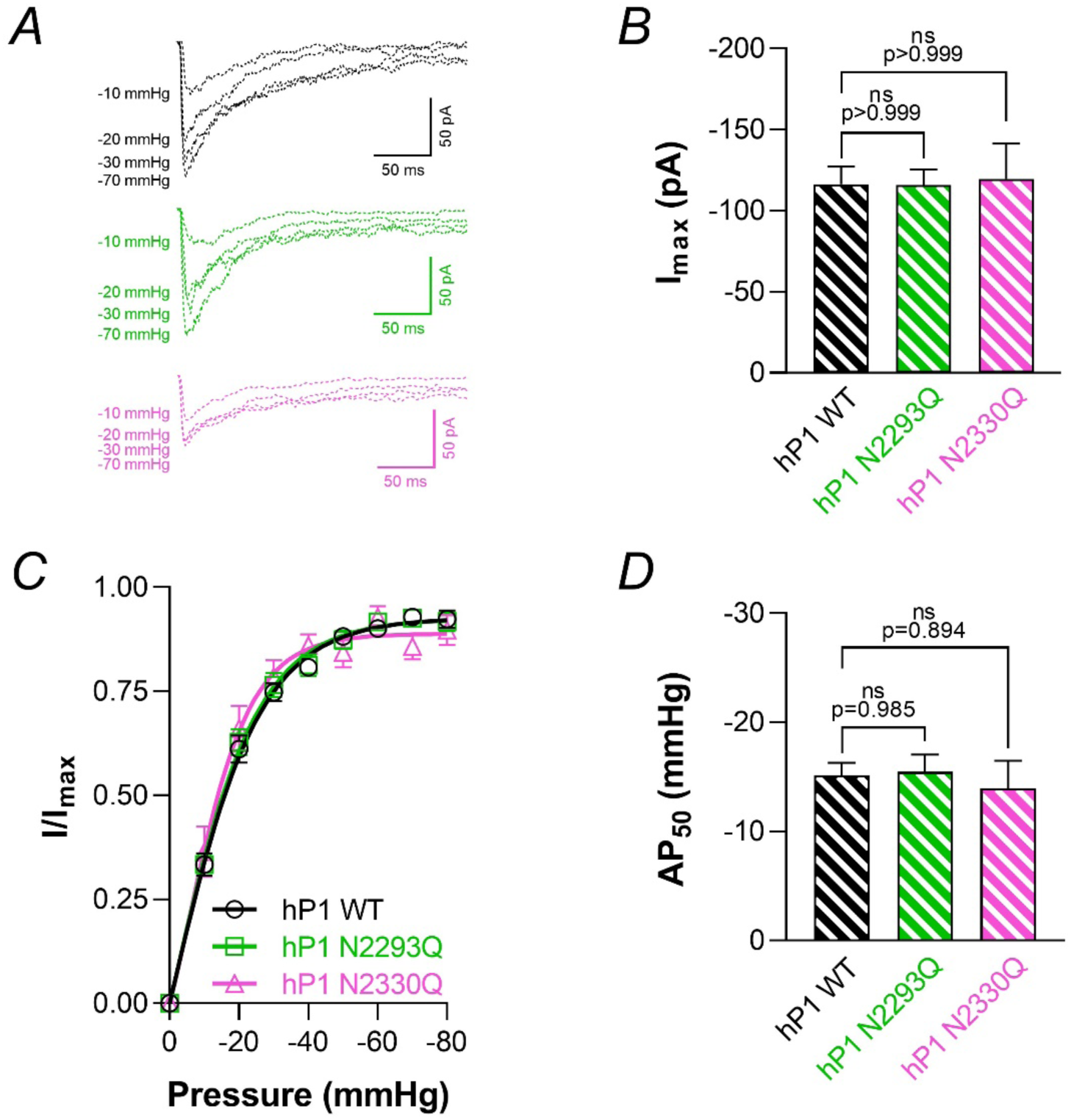
Piezo1 mechanosensitivity does not depend on its glycosylation status when using collagen-coated substrates. *A*, representative traces of mechanically activated currents obtained from cell-attached patches on HEK293 cells heterologously expressing the WT, N2293Q or N2330Q hPiezo1 channel, as indicated, using collagen as the surface coating agent. Only traces in response to −10, −20, −30 and −70 mmHg pulses are shown for visualisation purposes. *B*, quantification of the peak current per patch (I_max_) elicited by negative pressure pulses. *C*, normalised current-pressure relationship calculated as the I/I_max_ ratio. *D*, average half-maximal activating pressure (AP_50_) for WT, N2293Q or N2330Q hPiezo1 channels, as indicated. Data presented as mean ± SEM. No significant differences detected via Kruskal-Wallis test (ns: not significant; exact P values shown when available; sample sizes: WT, n = 34; N2993Q, n = 25; N2330Q, n = 13).

On PLL substrates, I_max_ for N2293Q (−129.9 ± 15.5 pA, n = 31) and N2330Q (−100.1 ± 7.62 pA, n = 27) remained comparable to WT (−157.0 ± 23.08 pA, n = 20) (P = 0.574 and P = 0.057, respectively) (Fig. 3*A* and *B*). Strikingly, both mutants exhibited significantly lower AP_50_ values (N2293Q: −11.72 ± 1.1 mmHg, n = 31; N2330Q: −11.05 ± 0.96 mmHg, n = 27) versus WT (−20.53 ± 2.26 mmHg, n = 20, P < 0.001), indicating heightened mechanosensitivity (Fig. 3*C* and *D*). Given these results, we focus the rest of our work on cells exposed to PLL as substrate.

**Figure 3.**
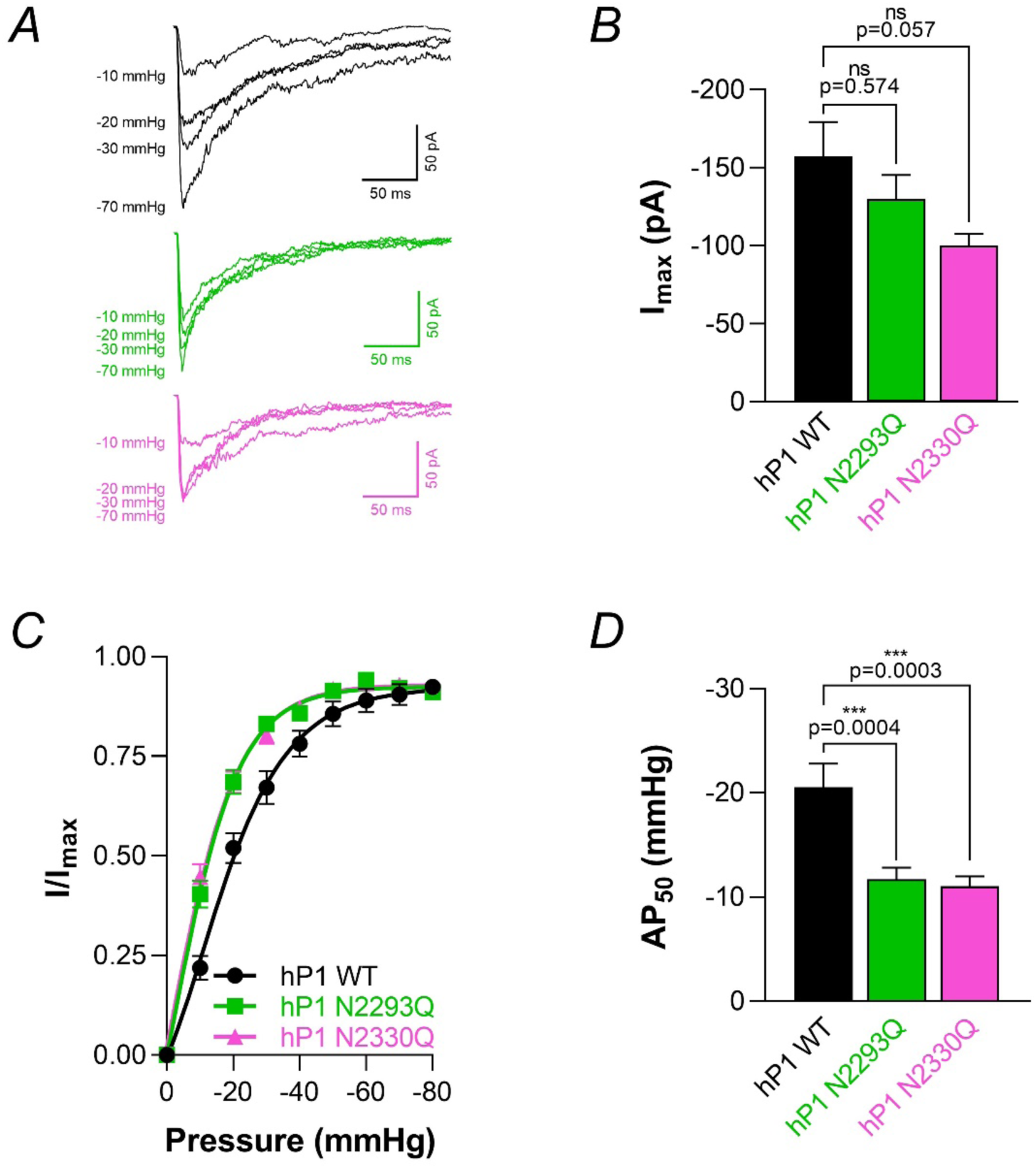
Piezo1 N-glycosylation mutants show an increased mechanical sensitivity on PLL-coated substrates. *A*, representative traces of mechanically activated currents obtained from cell-attached patches on HEK293 cells heterologously expressing the WT, N2293Q or N2330Q hPiezo1 channel, as indicated, using PLL as the surface coating agent. Only traces in response to −10, −20, −30 and −70 mmHg pulses are shown for visualisation purposes. *B*, quantification of the peak current per patch (I_max_) elicited by negative pressure pulses. *C*, normalised current-pressure relationship calculated as the I/I_max_ ratio. *D*, average half-maximal activating pressure (AP_50_) for WT, N2293Q or N2330Q hPiezo1 channels, as indicated. Data presented as mean ± SEM. Data presented as mean ± SEM. Statistical significance versus WT was assessed using Kruskal-Wallis test followed by Dunn’s post hoc test. (***P < 0.001; ns, not significant; exact P values shown; sample sizes: WT, n = 20; N2993Q, n = 31; N2330Q, n = 27).

Inactivation time constants (τ_inact_) for N2293Q and N2330Q mutants on PLL did not differ from WT (P > 0.999) (Fig. 5), consistent with prior reports that single-site N-glycosylation modifications in the cap domain do not alter Piezo1 inactivation kinetics (Li, Ng et al., 2021).

### Pharmacological Recapitulation of PMM2-CDG Hypoglycosylation Reveals Mature N-Glycans as Critical Determinants of Piezo1 Mechanical Force Sensing

So far, our findings align with established glycobiology methodologies using asparagine mutagenesis to probe glycosylation-dependent functions (Brunner et al., 1992; Sandoval et al., 2004; Tétreault et al., 2016; Izquierdo-Serra et al., 2018; Virion et al., 2019; Li, Ng et al., 2021). This conventional experimental approach of inducing complete N-glycosylation ablation at specific sites fails to adequately replicate the pathophysiological continuum characteristic of conditions like Phosphomannomutase 2-Congenital Disorder of Glycosylation (PMM2-CDG), where residual enzymatic activity allows for limited but dysfunctional glycan processing. To better model this clinically relevant hypoglycosylation paradigm, we implemented pharmacological inhibition of N-glycan maturation through established mechanisms (Tulsiani & Touster, 1983; Elbein et al., 1990). We employed swainsonine (Sw; 250 µM) and kifunensine (Kf; 25 µM), thereby creating a more physiologically relevant experimental system for studying the spectrum of glycosylation deficiencies observed in human glycoprotein disorders. Under these experimental conditions we observed a clear hypoglycosylation of WT Piezo1 (Fig. 4). Neither inhibitor altered WT Piezo1 current amplitude (Sw: −195.1 ± 24.53 pA, n = 31; Kf: −130.1 ± 15.97 pA, n = 7; vs. control: −157.0 ± 23.08 pA, n = 20; Kruskal-Wallis P ≥ 0.446; Fig. 6*A* and *B*) or inactivation kinetics (P ≥ 0.659) (Fig. 5) in cell attached-patches from HEK293 cells expressing the channel exposed to negative pressure pulses. However, both agents significantly reduced AP_50_ values (Sw: −11.29 ± 1.33 mmHg, n = 18; Kf: −10.93 ± 1.3 mmHg, n = 7; vs. control: −20.53 ± 2.26 mmHg; Kruskal-Wallis/Dunn’s test P < 0.01 and P < 0.05, respectively; Fig. 6*C* and *D*), mirroring the hypersensitivity phenotype of N2293Q and N2330Q mutants.

**Figure 4.**
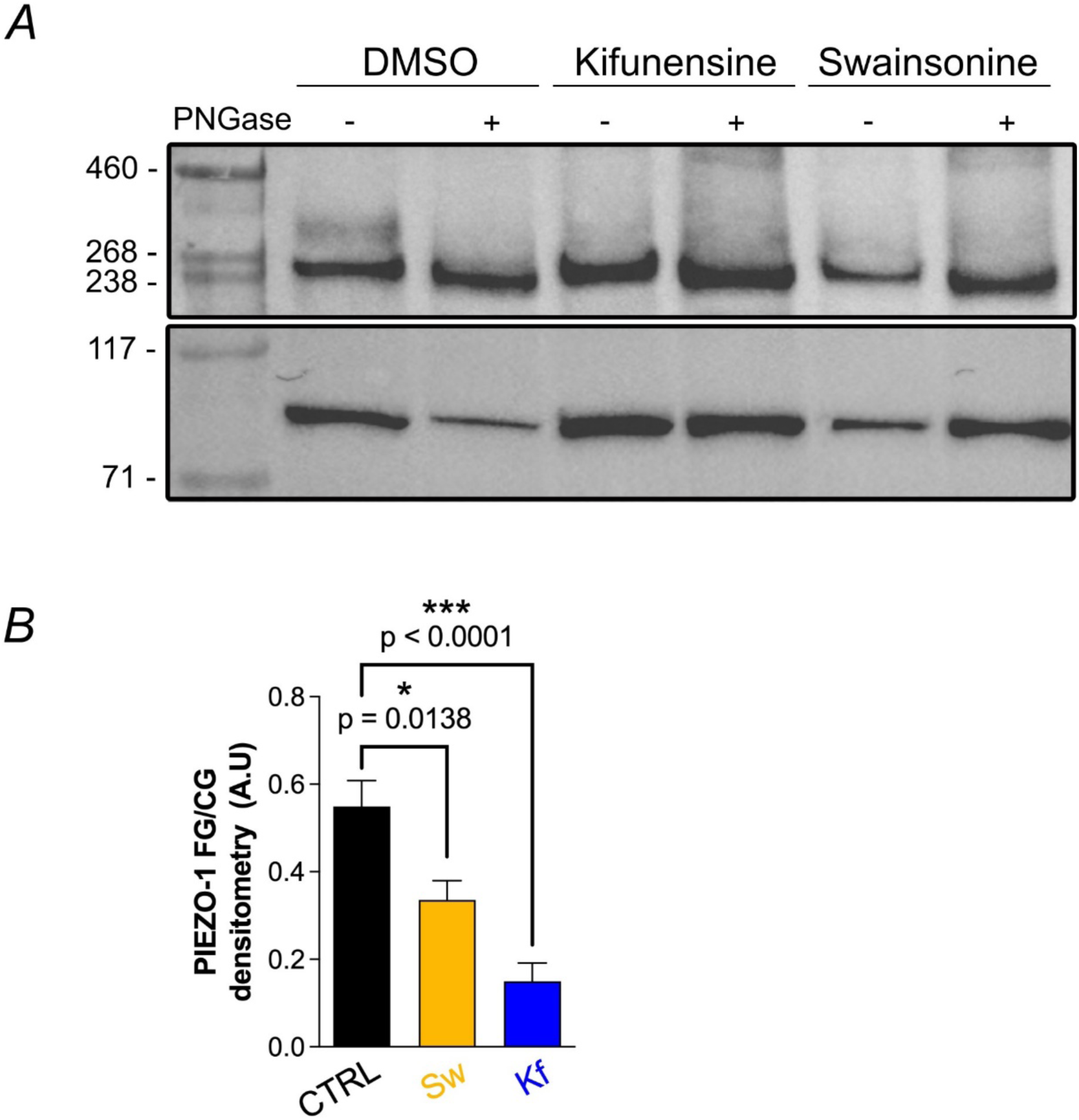
Chemically-induced inhibition of WT Piezo1 mature glycosylation leads to lower level of channel full glycosylation. *A*, Representative blot showing the effect on heterologously expressed WT Piezo1 glycosylation in HEK293 cells treated with DMSO (1:1000), 250 μM swainsonine, or 25 μM kifunensine for 48 hours. PNGase F was also added to protein extraction from each experimental condition to identify the hypoglycosylated form of WT Piezo1 in vitro (corresponding to the lower core-glycosylated (CG) band induced by cleavage of the upper fully-glycosylated (FG) band of the channel, as previously reported (Li et al., 2021)). *B*, Quantification of the Piezo1 FG/CG ratio after control treatment (CTRL, DMSO (1:1000) and in the presence of the two inhibitors of mature glycosylation. ***P < 0.0001 and * P < 0.05 determined by one-way ANOVA with Dunnett’s post-hoc test; exact P value shown when available.

**Figure 5.**
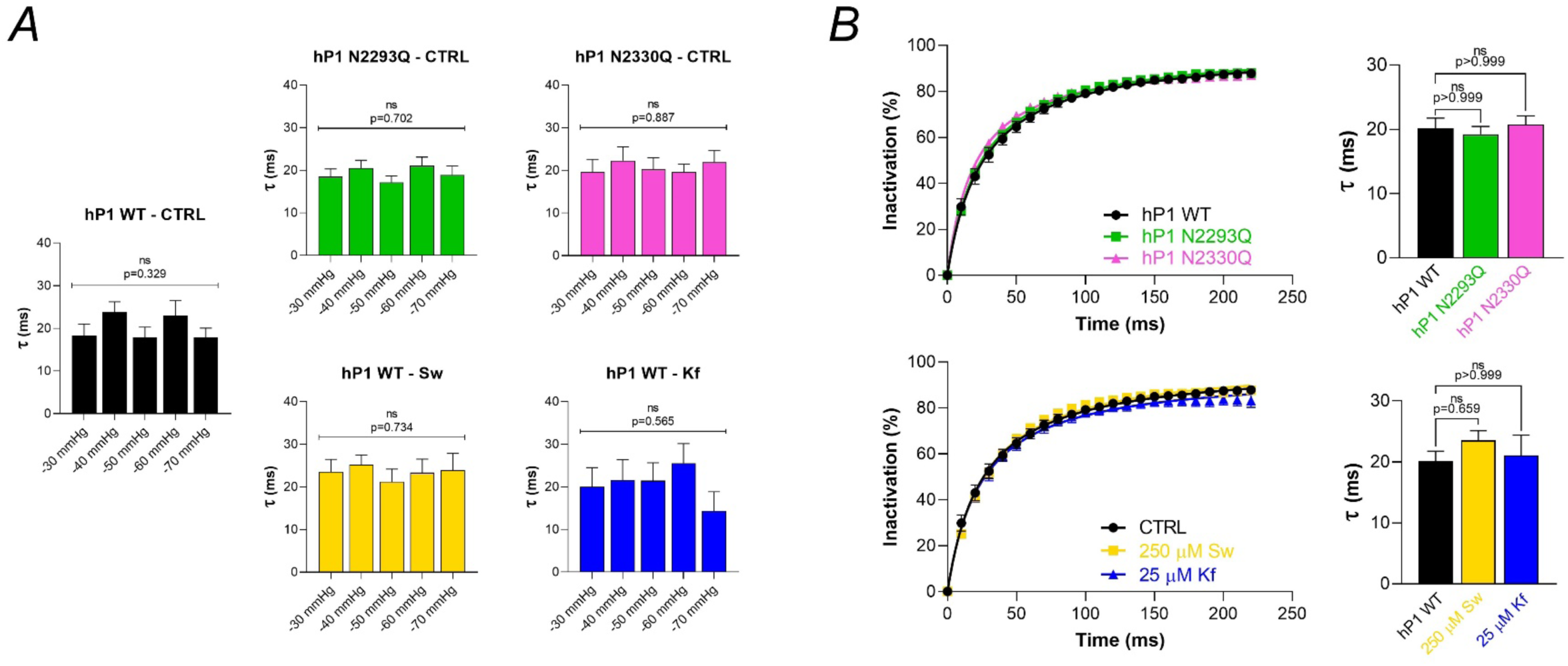
Piezo1 hypoglycosylation does not affect its inactivation kinetics. *A*, time constant of inactivation (τ) of mechanically activated currents obtained from cell-attached patches on HEK293 cells overexpressing the WT or the corresponding N-glycosylation hPiezo1 mutants, as indicated. When using the glycosylation inhibitors swainsonine (Sw) or kifunensine (Kf), cells were treated for 48 hours. *B*, inactivation curve showing the percentage of inactivation from the peak current at the indicated time points of the applied negative pressure pulse (left), and time constant of inactivation (τ) (right). Data were averaged for pulses from −30 mmHg through −70 mmHg. Data presented as mean ± SEM. No significant differences detected via Kruskal-Wallis test (ns: not significant; exact P values shown).

**Figure 6.**
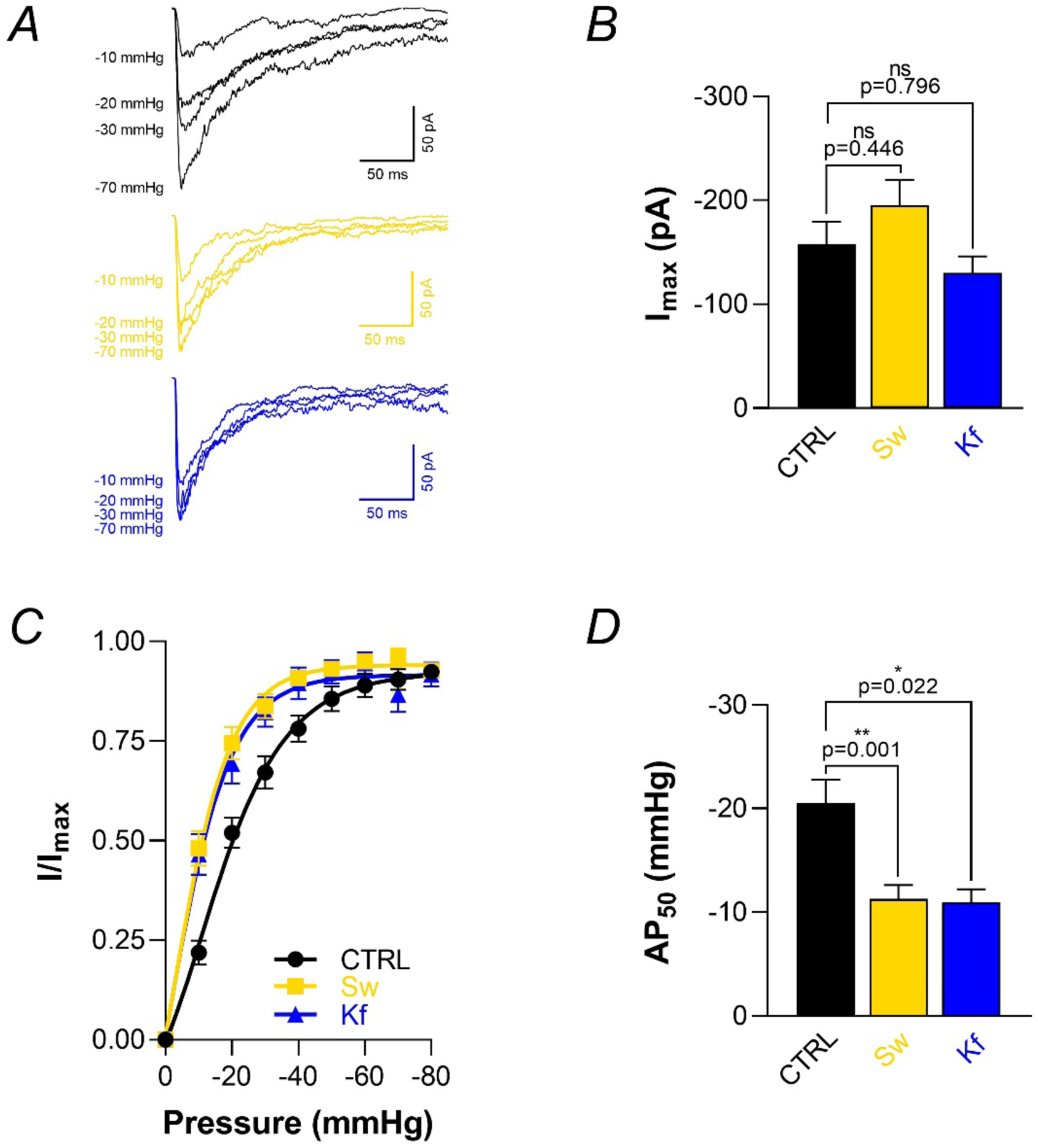
Chemically induced inhibition of WT Piezo1 mature glycosylation leads to an increased mechanosensitivity. *A*, representative traces of mechanically activated currents obtained from cell-attached patches on HEK293 cells heterologously expressing the WT hPiezo1 channel and treated for 48 hours with 250 µM swainsonine or 25 µM kifunensine, as indicated. Recordings were performed using PLL as the surface coating agent. Only traces in response to −10, −20, −30 and −70 mmHg pulses are shown for visualisation purposes. B, quantification of the peak current per patch (I_max_) elicited by negative pressure pulses. C, normalised current-pressure relationship calculated as the I/I_max_ ratio. D, average half-maximal activating pressure (AP_50_) for each experimental condition. Data presented as mean ± SEM. Statistical significance versus vehicle control (CTRL) was assessed using Kruskal-Wallis test followed by Dunn’s post hoc test. (*P < 0.05; **P < 0.01; ns, not significant; exact P values shown; sample sizes: CTRL, n = 20; Sw, n = 18; Kf, n = 7).

The concordance between mutagenesis and pharmacological inhibition demonstrates that disrupted N-glycan maturation, and not merely glycan absence, underlies enhanced mechanosensitivity. This suggests Piezo1’s glycan-dependent mechanical tuning requires mature glycan structures, with immature intermediates (e.g., hybrid glycans in Sw-treated cells) insufficient to maintain WT-like force transduction.

### Hypoglycosylation Lowers Piezo1’s Mechanical Activation Threshold to Potentiate Calcium Influx Under Submaximal Cellular Stretch

Our cell-attached patch-clamp recordings in HEK293 cells enabled direct electrophysiological characterisation of Piezo1 mechanosensitivity under hypoglycosylation conditions. To complement our findings and assess channel activation within an integrated cellular framework, we employed calcium imaging coupled with controlled uniaxial stretch stimulation, thereby evaluating mechanically induced Piezo1 activity across physiological contexts.

Sequential 40% and 80% uniaxial stretch challenges (10-minute interval) elicited significant calcium influx in HEK293 cells expressing WT Piezo1 only at the higher stretch intensity (80% stretch). In contrast, cells expressing N2293Q or N2330Q Piezo1 mutants exhibited enhanced calcium influx at 40% stretch compared with cells expressing the WT channel (Fig. 7*A*-*C*). This enhancement was particularly consistent and significant for the N2993Q mutant when measuring both the peak amplitude and the area-under-curve (AUC) of the stretching-induced transient Ca^2+^ signal (Kruskal-Wallis/Dunn’s test P < 0.0001 and P = 0.0003, respectively). At higher mechanical stimuli (80% stretch), Ca^2+^ responses mostly converged between cells expressing mutant and WT channels (Fig. 7*A*-*C*), mirroring the loss of differential sensitivity observed in cell-attached patch-clamp recordings exposed to pressures exceeding −40 mmHg (Fig. 3*C*).

**Figure 7.**
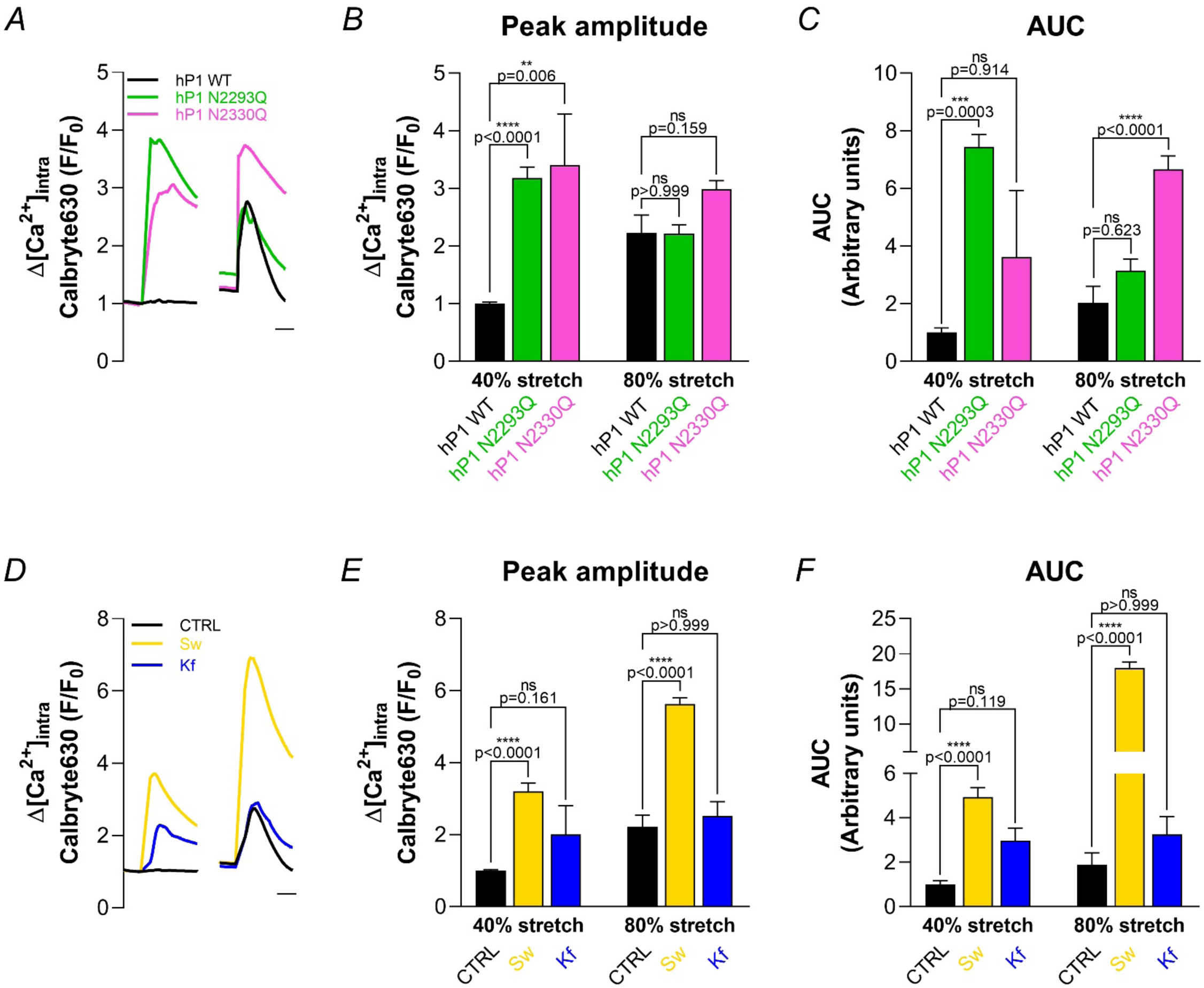
Ca^2+^ influx trough Piezo1 channels under submaximal stretch is enhanced by hypoglycosylation. *A* and *D*, mean intracellular Ca^2+^ signals obtained from HEK293 cells transfected with WT, N2293Q or N2330Q hPiezo1 (as indicated) (Calbryte630 AM loading). Treatments for 48 hours with 250 µM swainsonine (Sw) or 25 µM kifunensine (Kf) were applied when indicated. Arrows denote 40% and 80% uniaxial stretch (10-minute interval). Scale bar: 1 minute. *B* and *E*, normalised peak Ca^2+^ transient amplitudes following mechanical stretch. *C* and *F*, quantification of integrated Ca^2+^ responses (area under curve, AUC) for stretch-induced signals. Data presented as mean ± SEM (numbers in bars indicate cell counts pooled from 3 biological replicates). Statistical significance versus WT or vehicle control (CTRL) was assessed using Kruskal-Wallis test followed by Dunn’s post hoc test. (**P < 0.01; ***P < 0.001; ****P < 0.0001; ns, not significant; exact P values shown when available).

Pharmacological disruption of glycosylation maturation using swainsonine (Sw, 250 µM) or kifunensine (Kf, 25 µM) recapitulated this potentiated mechanosensitive phenotype. Swainsonine enhanced Ca^2+^ responses to both 40% and 80% uniaxial stretch pulses (Fig. 7*D*-*F*). This contrasted with kifunensine treatment, which showed a non-significant tendency to increase responses only at 40% stretch, suggesting distinct roles for complex versus hybrid/high-mannose glycans in regulating Piezo1 channel function.

These results demonstrate that Piezo1 hypoglycosylation potentiates mechanosensitivity at the cellular level, increasing calcium influx following submaximal uniaxial stretch. The concordance between electrophysiological and calcium imaging data establishes glycosylation as a critical modulator of Piezo1’s mechanical activation threshold across multiple experimental paradigms.

### Hypoglycosylation Induces Upregulation of Somatic Piezo1 Expression While Preserving Synaptic Localisation Patterns in Murine Cortical Neurons

HEK293 cells provide technical advantages for electrophysiological characterisation of heterologously expressed ion channels, such as ease of transfection, rapid growth, and robust protein expression, which collectively enable high-throughput screening and detailed biophysical analyses (Thomas & Smart, 2005). However, HEK293 cells often fail to fully recapitulate the native cellular environment of the protein under investigation, including its regulatory mechanisms, a critical consideration when studying complex disorders like CDGs. Consequently, Piezo1’s cellular context, post-translational modifications, and interacting partners in HEK293 cells may diverge from those in native tissues, potentially limiting the physiological relevance of findings (Hunter et al., 2019; Zhou et al., 2023). Murine cortical neurons, which endogenously express mechanosensitive channels compatible with Piezo1 (Coste et al., 2010; Nikolaev et al., 2015), offer more representative model for investigating neurological manifestations that are a hallmark of CDGs (Serrano, 2021; Freeze et al., 2015; Francisco et al., 2023).

We first confirmed Piezo1 expression in primary murine cortical neurons. Confocal imaging of neurons immunostained against Piezo1 revealed robust channel distribution across somatic and neuritic compartments (Fig. 8).

**Figure 8.**
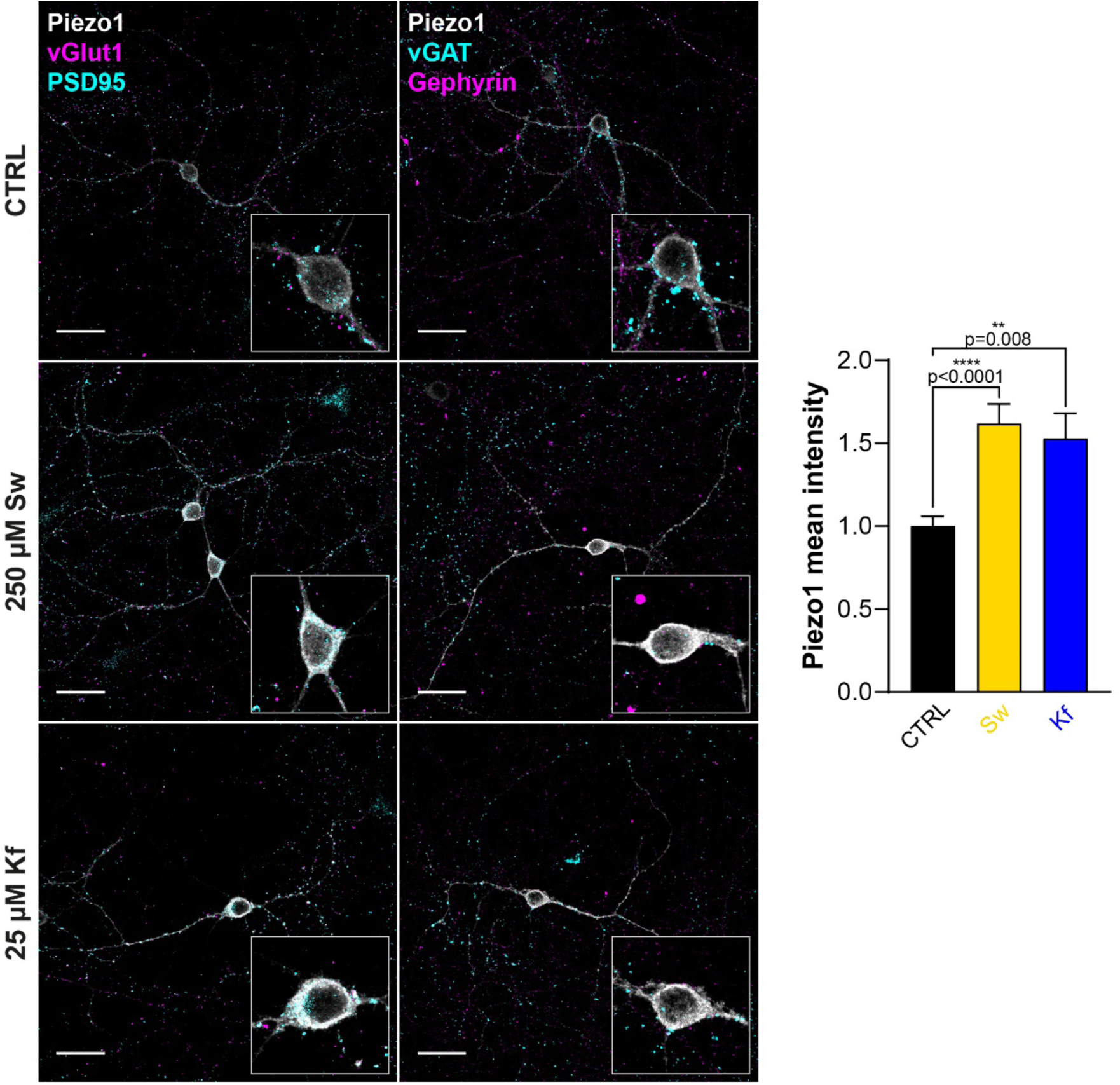
Swainsonine and kifunensine increase somatic Piezo1 levels in murine cortical neurons. *A*, confocal images of primary cortical neurons from mice co-immunostained for Piezo1 and excitatory (vGlut1/PSD95) or inhibitory (vGAT/Gephyrin) synaptic markers. Neurons were treated with 250 µM swainsonine (Sw) or 25 µM kifunensine (Kf) for 72 hours. Scale bar: 25 µm. Inset: somatic region magnified. *B*, quantification of Piezo1 mean intensity at the soma. Data presented as mean ± SEM. Statistical significance versus vehicle control (CTRL) was assessed using Kruskal-Wallis test followed by Dunn’s post hoc test. (**P < 0.01; ****P < 0.0001; sample sizes: CTRL, n = 43; Sw, n = 35; Kf, n = 27 (n = neurons pooled from 5 biological replicates)).

To assess whether hypoglycosylation alters Piezo1 neuronal expression or synaptic localisation, we performed dual immunostaining against Piezo1 and presynaptic/postsynaptic markers (vGlut1/PSD95 for excitatory synapses; vGAT/Gephyrin for inhibitory synapses). Swainsonine and kifunensine treatments increased somatic Piezo1 levels by 62 ± 12% (n = 35, P < 0.0001) and 53 ± 15% (n = 27, P = 0.008), respectively (Fig. 8). However, co-localisation analyses using the Manders Overlap Coefficient (MOC) demonstrated no significant alteration in Piezo1’s synaptic localisation at excitatory or inhibitory synapses (Fig. 9). Furthermore, quantification of Piezo1 fluorescence intensity at synaptic puncta (vGlut1, vGAT, PSD95, or Gephyrin) revealed no changes following N-glycosylation inhibition (Fig. 10).

**Figure 9.**
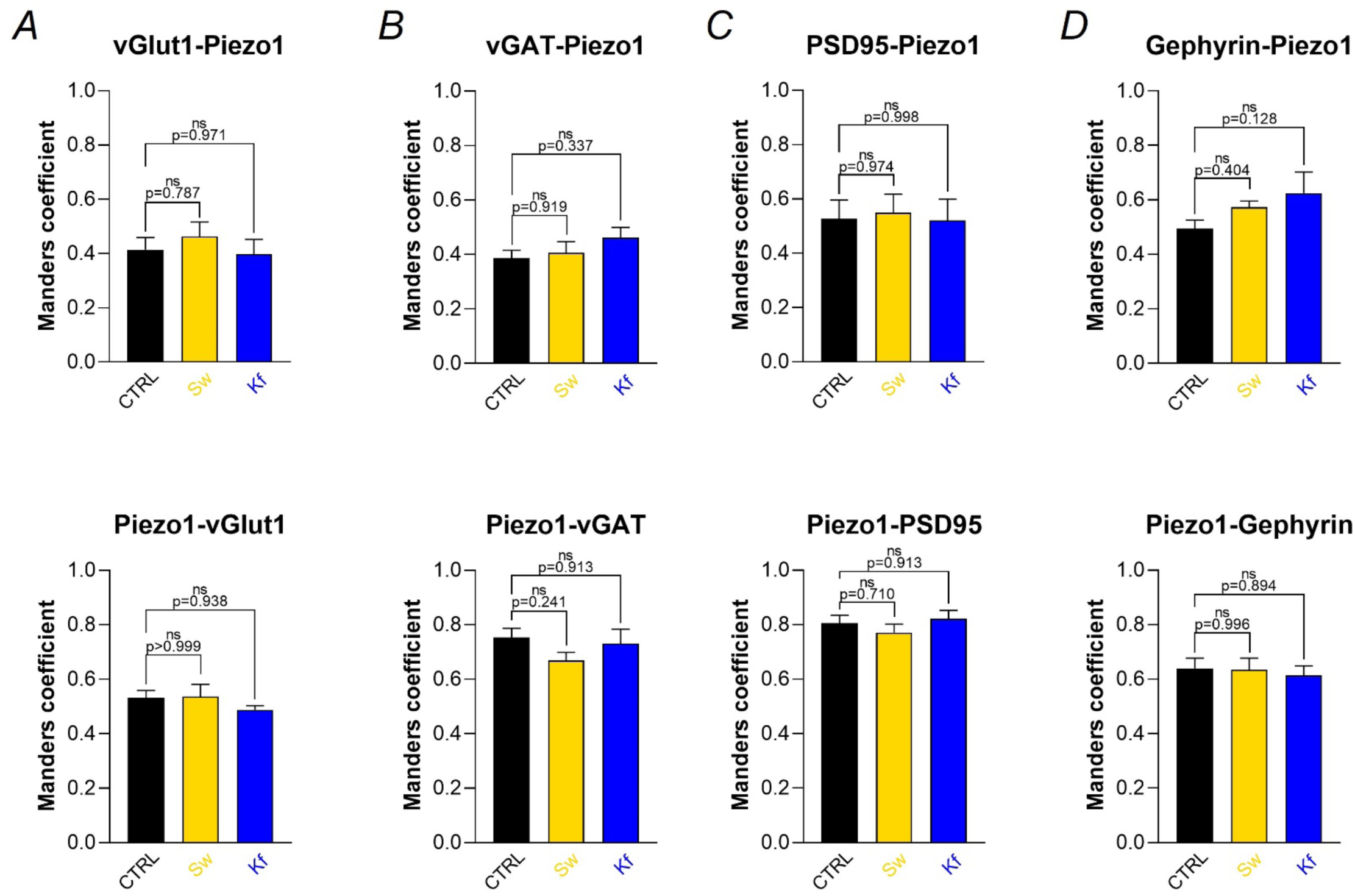
Swainsonine and kifunensine preserve Piezo1 synaptic localisation patterns. *A-D*, co-localisation of Piezo1 with the presynaptic/postsynaptic markers, quantified via the Manders Coefficient. Data presented as mean ± SEM (numbers in bars indicate cell counts pooled from 4 biological replicates). No significant differences detected via Kruskal-Wallis test (ns: not significant; exact P values shown).

**Figure 10.**
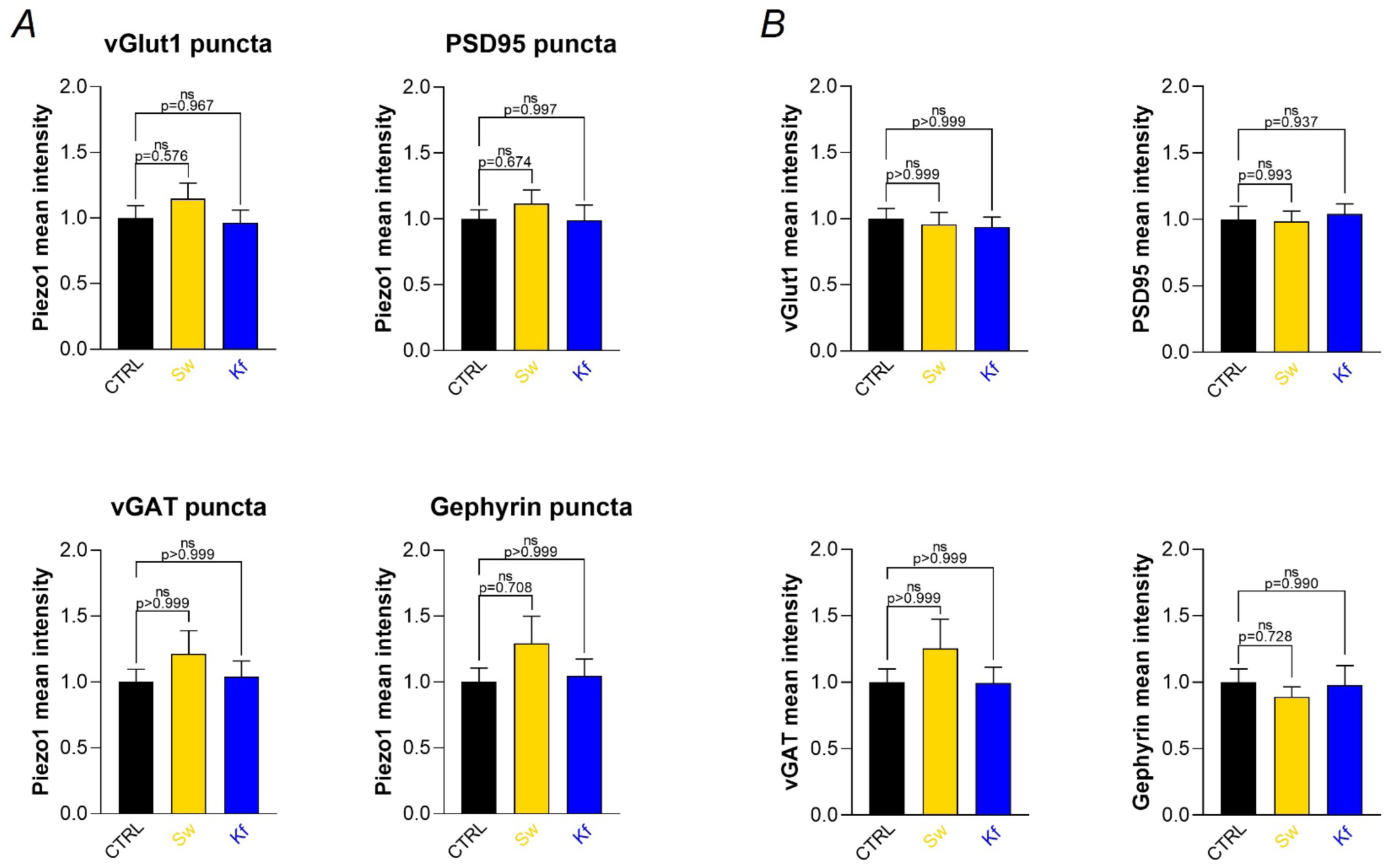
Swainsonine and kifunensine do not alter Piezo1 synaptic levels. *A*, Piezo1 intensity at synaptic puncta marked by vGlut1, vGAT, PSD95 or Gephyrin. *B*, synaptic markers intensity within excitatory (vGlut1/PSD95) and inhibitory (vGAT/Gephyrin) synapses. Data presented as mean ± SEM (numbers in bars indicate cell counts pooled from 4 biological replicates). No significant differences detected via Kruskal-Wallis test (ns: not significant; exact P values shown when available).

### Hypoglycosylation Enhances Piezo1-Mediated Calcium Responses in Murine Cortical Neurons

Following the characterisation of Piezo1 expression patterns in murine primary cortical neurons, we examined how pharmacological manipulation of N-linked glycosylation impacts channel functionality. Comparative calcium imaging between neurons treated for 72 hours with swainsonine (250 μM), kifunensine (25 μM), or vehicle controls demonstrated significant potentiation of Piezo1-dependent calcium responses through two distinct activation modalities.

Application of the Piezo1-specific activator Yoda1 (10 μM, 30s) elicited markedly enhanced calcium influx in treated neurons, with both peak amplitude (swainsonine: 1.94 ± 0.24-fold increase, n = 41, P < 0.0001 *vs* control; kifunensine: 1.95 ± 0.16-fold increase, n = 7, P < 0.01 *vs* control) and integrated signal (AUC) (swainsonine 2.99 ± 0.35-fold increase, P < 0.0001 *vs* control; kifunensine 2.76 ± 0.47-fold increase, P < 0.05 *vs* control) demonstrating robust potentiation (Fig. 11*A*-*C*). This enhancement of the Ca^2+^ response to Yoda1 extended to mechanical stimulation, where 40% uniaxial stretch pulses produced 1.91 ± 0.29-fold increase (swainsonine, n=12) and 3.27 ± 0.58-fold increase (kifunensine, n=23) increases in peak Ca^2+^ transient amplitude compared to controls (P < 0.05 and P < 0.001 respectively; Fig. 11*D*-*E*). Notably, this potentiation of the Ca^2+^ signal amplitude exhibited stimulus intensity dependence, as 80% stretch pulses failed to elicit differential responses between treatment groups (Fig. 11*D*-*E*), mirroring our previous results in HEK293 cells that suggest hypoglycosylation preferentially enhances submaximal mechanical activation of Piezo1. Signal integral analysis revealed treatment-specific potentiation patterns, with kifunensine showing significant enhancement at 40% stretch (P < 0.001) and swainsonine demonstrating augmented responses at 80% stretch (P < 0.001) (Fig. 11*F*).

**Figure 11.**
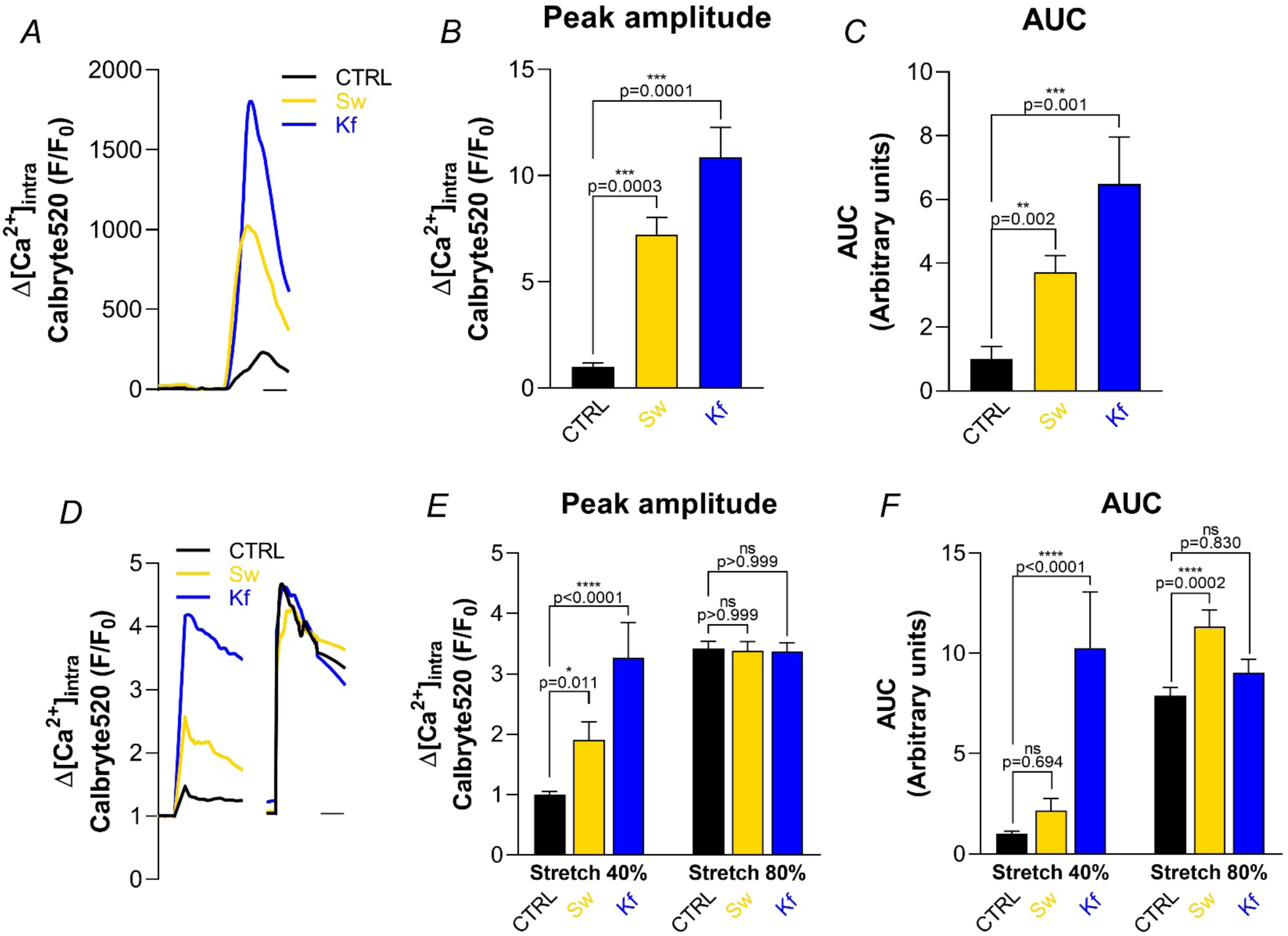
Pharmacological inhibition of glycan maturation enhances Yoda1- and stretch-induced Ca^2+^ responses in murine cortical neurons. *A* and *D*, mean intracellular Ca^2+^ signals from Calbryte520 AM-loaded primary murine cortical neurons treated for 72 hours with swainsonine (Sw, 250 µM) or kifunensine (Kf, 25 µM). Traces show responses to either: 10 µM Yoda1 perfusion (30 s pulse) (*A*), or sequential 40% and 80% uniaxial stretch pulses (10 min interstimulus interval) (*D*). Scale bar: 1 min. *B* and *E*, normalised peak Ca^2+^ transient amplitudes following Yoda1 application (*B*) or mechanical stretch (*E*). *C* and *F*, quantification of integrated Ca^2+^ responses (area under curve, AUC) for Yoda1-evoked (*C*) and stretch-induced (*F*) signals. Data presented as mean ± SEM (numbers in bars indicate cell counts pooled from 3 biological replicates). Statistical significance versus vehicle control (CTRL) was assessed using Kruskal-Wallis test followed by Dunn’s post hoc test (*P < 0.05, **P < 0.01, ***P < 0.001, ****P < 0.0001; ns = not significant; exact P values shown when available).

The conserved pattern of hypoglycosylation-induced potentiation across both pharmacological and mechanical activation modalities suggests a generalised enhancement of Piezo1 function through N-linked glycan modification in neuronal systems.

## Discussion

### Mature N-glycans set the mechanical activation threshold of Piezo1

The principal finding of this study is that mature N-linked glycans act as determinants of the force threshold for Piezo1 activation. Three complementary observations support this conclusion. First, removal of either of the two conserved cap-domain N-glycosylation sites shifted the pressure-response relationship of Piezo1 towards lower activating pressures on poly-L-lysine (PLL). Second, two pharmacological interventions that restrict N-glycan maturation, swainsonine or kifunensine, reproduced this shift in wild-type Piezo1. Third, the electrophysiological phenotype was evident at the cellular level as greater Ca^2+^ entry during submaximal stretch and, in primary mouse cortical neurons, as enhanced responses to both mechanical stimulation and the Piezo1 activator Yoda1. Importantly, hypoglycosylation did not increase maximal patch current or slow inactivation. The effect is therefore best interpreted as sensitisation of channel activation rather than an increase in maximal current-carrying capacity or prolongation of the open state.

Glycosylation sites N2294 and N2331 in the human Piezo1 cap domain (N2293 and N2330 in the isoform used here) were previously shown to be major determinants of channel maturation and trafficking. Thus, simultaneous removal of both sites severely impairs surface expression, whereas either N-glycosylation site alone is sufficient to sustain measurable mechanically activated currents (Li, Ng et al., 2021). The present results extend that study by showing that channels retaining functional expression after loss of one N-glycosylation site are not biophysically equivalent to the fully glycosylated channel. Similar peak currents across genotypes indicate that the number of functional channels captured in the patch, their unitary conductance, and their maximal open probability were not grossly altered under the present conditions. By contrast, the marked decrease in half-maximal activating pressure indicates that glycan-deficient channels require less mechanical input to enter conducting states. Thus, the two cap-domain N-glycosylation sites appear to make partly separable contributions to Piezo1 biology, as retention of either site is sufficient to support productive trafficking, whereas preservation of the normal mechanical activation range requires both sites to be glycosylated and depends more broadly on proper N-glycan maturation.

### Mechanistic basis of glycosylation-dependent Piezo1 Gating

Piezo1 mechanosensitivity is regulated by cytoskeletal organisation, adhesion molecules, and the composition of the extracellular matrix (ECM), among other factors. Channel activation has been associated with F-actin thickening and focal adhesion enlargement (Jetta et al., 2023; Morena et al., 2024), while interaction of the E-cadherin ectodomain with the central cap increases the I_max_, reduce the AP_50_ and increase the open probability of Piezo1 (Wang et al., 2022). In addition, Piezo1 exhibits increased mechanosensitivity on collagen than on PLL, both in atomic force microscopy assays (Gaub & Müller, 2017) and under shear stress (Lai et al., 2022). Consistent with these observations, wild-type Piezo1 in our hands was likewise more mechanosensitive on collagen-coated substrates.

The extracellular cap is a plausible structural substrate for these N-glycan effects. Piezo1 is a trimeric, curved membrane protein in which peripheral blades and intracellular beams transmit force over long distances to a central pore capped by a large extracellular domain (Saotome et al., 2018). This cap contributes not only to pore architecture but also to mechanical gating and inactivation (Lewis & Grandl, 2020). Because the two functionally relevant N-glycosylation sites lie within this cap, they are well positioned to influence the energetic coupling between force-sensing elements and pore opening through steric constraints, hydration, electrostatic interactions, or effects on cap mobility and conformational flexibility. Indeed, crosslinking cap subdomains to constrain their conformational flexibility abolishes Piezo1 mechanical activation (Lewis & Grandl, 2020). In this context, glycosylation of the central cap may influence Piezo1 mechanosensitivity by modulating the domain’s conformational flexibility, where the absence of mature glycans increases cap flexibility and thereby reduces the mechanical energy required for conformational transitions leading to pore opening. This parallels observations in the Piezo1 anchor domain, where glycine insertions at residue P2113 enhanced mechanosensitivity by disrupting interactions between the anchor and inner helices, increasing conformational flexibility (Li, Cox et al., 2021). The adjacent F2114 residue, which is critical for coupling mechanical forces to pore opening, may adopt altered conformations in hypoglycosylated Piezo1, further reducing the activation threshold by removing steric hindrance on the inner pore-lining helix (Li, Cox et al., 2021). Cap glycosylation may also modulate channel mechanosensitivity indirectly, by influencing interactions with extracellular partners such as E-cadherin (Wang et al., 2022).

Evidence addressing the role of glycosylation in mechanosensitive ion channels remains limited, and where it exists, it points in different directions. Piezo1 expressed in N-acetylglucosaminyl-transferase I-deficient (GnT1^-/-^) HEK293 cells shows reduced mechanosensitivity in response to negative pressure in cell-attached recordings (Li, Ng et al., 2021). Because GnT1 is required for the transition from high-mannose to hybrid and complex N-glycans (Kornfeld & Kornfeld, 1985), these cells produce only high-mannose structures, similar to kifunensine-treated conditions. However, our data contrast with these findings, as both pharmacological inhibition of glycan maturation (swainsonine or kifunensine) and glycosylation site-directed single mutations (N2293Q, N2330Q) increased Piezo1 mechanosensitivity. The concordance between mutagenesis and pharmacological approaches supports a genuine glycan-dependent mechanism and indicates that proper glycan maturation, rather than mere occupancy of asparagine residues, is required for normal force sensing. Thus, our findings suggest that complex N-glycans normally dampen Piezo1’s baseline mechanosensitivity on low-adhesive surfaces such as PLL, possibly by stabilising the channel’s closed state. Conversely, under similar substrate conditions, the presence of either high-mannose or hybrid-type N-glycans is sufficient to enhance the channel’s mechanical sensitivity. A related principle applies to other mechanoreceptors. Indeed, β_2_ adrenergic receptors (β_2_AR) or ENaC rely on glycan tethers that transduce mechanical force from the ECM to the channel, and hypoglycosylation impairs their mechanical activation (Virion et al., 2019; Knoepp et al., 2020). Because hypoglycosylation has the opposite effect on Piezo1, glycans in the cap region likely act through a different, more intrinsic structural mechanism rather than as external tethers, a distinction that future comparisons of Piezo1 structures at defined glycosylation states, together with analysis of ECM-interaction changes, should help clarify. Identifying the precise structural basis will also require site-specific glycoproteomics and glycoengineered systems.

A notable finding is that the effect of hypoglycosylation depends on the adhesive substrate. Wild-type Piezo1 is more mechanosensitive on collagen than on PLL, consistent with the role of ECM composition and integrin engagement in force transmission. On collagen, the activation threshold of wild-type Piezo1 was already comparable to that of glycosylation-deficient channels, and the N2293Q and N2330Q mutations produced no further sensitisation. On PLL, by contrast, both mutations strongly enhanced mechanosensitivity. One interpretation is that collagen-dependent force transmission places the channel in an already mechanically primed state that occludes any further effect of glycan removal. Alternatively, glycans may participate in functionally relevant extracellular interactions that are altered when integrin-mediated coupling is weak or qualitatively different, as expected on PLL-coated substrates (Toyoshima & Nishida, 2007). More broadly, Piezo1 responds to bilayer tension but is also shaped in intact cells by the actin cytoskeleton and cadherin-catenin complexes (Wang et al., 2022); N-glycans could modulate either route, exerting steric or entropic effects on the cap itself while also influencing adhesion-receptor conformation, clustering, and membrane-cortex coupling. Piezo1 mechanosensitivity thus likely emerges from an interplay between intrinsic channel properties and the mechanical architecture of the cell surface.

Finally, hypoglycosylation left inactivation kinetics unchanged, even though subdomains at the base of the cap strongly influence inactivation (Lewis & Grandl, 2020) and gain-of-function variants associated with dehydrated hereditary stomatocytosis prolong channel activity by slowing it (Zarychanski et al., 2012; Demolombe et al., 2013). This is consistent with previous reports showing that N2293Q and N2330Q single mutants inactivate similarly to wild-type Piezo1 (Li, Ng et al., 2021). At the mechanistic level, these observations indicate that hypoglycosylation in the cap region of Piezo1 produces a different gain-of-function phenotype that is independent of channel inactivation and instead characterised by increased recruitment of Piezo1 by weak or intermediate mechanical stimuli. This argues against a nonspecific destabilisation of channel gating and supports a model in which glycan trimming selectively affects force-dependent gating rather than the intrinsic biophysical properties of the channel.

### Hypoglycosylation expands the range of mechanical stimuli that evoke Ca^2+^ entry

The stretch experiments clarify the functional meaning of the pressure-response shift. In transfected HEK293 cells, glycosylation-deficient Piezo1 channels generated substantially larger Ca^2+^ transients at 40% stretch, whereas responses generally converged at 80% stretch. A comparable stimulus dependence was observed in cortical neurons treated with the glycosylation inhibitors swainsonine or kifunensine. Mechanical stimuli that are subthreshold, or only weakly effective, for fully glycosylated Piezo1 thus become sufficient to elicit appreciable cation entry after hypoglycosylation, consistent with a lowered activation threshold. At stronger mechanical stimulation, recruitment of the remaining channel population narrows the difference between conditions. This threshold-centred interpretation is consistent with the unchanged maximal patch currents and inactivation kinetics described above.

Enhanced Yoda1 responses in cortical neurons point to a second, complementary component of the phenotype. Yoda1 acts directly on Piezo1 and lowers its mechanical activation energy (Syeda et al., 2015), whereas stretch can also recruit other mechanosensitive ion channels expressed in cortical neurons, including TRPV1, TRPV2, TRPV4, TRPC1, TRPM7 and the TRPP1/2 complex (Christensen & Corey, 2007; Ranade et al., 2015; Yoo et al., 2022). With the exception of TRPC1, all of these channels have been reported to undergo N-glycosylation (Wirkner et al., 2005; Xu et al., 2006; Perálvarez-Marín et al., 2013; Hofherr et al., 2014; Overton et al., 2015), so their contribution to the enhanced stretch-evoked Ca^2+^ influx in cortical neurons cannot be excluded. The roughly twofold increase in Yoda1-evoked peak Ca^2+^ signals seen with either glycosylation inhibitor is more directly attributable to Piezo1 itself, and could reflect increased somatic Piezo1 abundance, altered allosteric coupling between the Yoda1-sensitive region and the pore, and/or amplification by downstream Ca^2+^ signalling. The parallel rise in somatic Piezo1 immunoreactivity and the gain-of-function phenotype observed in transfected HEK293 cells together support a direct contribution of Piezo1 hypoglycosylation to the increased stretch-induced Ca^2+^ responses in primary cortical neurons.

The absence of detectable changes in Piezo1 overlap with vGlut1, PSD95, vGAT, or gephyrin argues against a broad redistribution of Piezo1 towards excitatory or inhibitory synapses upon impaired N-glycan maturation, and against a simple increase in synaptic channel abundance as the basis for the enhanced Ca^2+^ responses. The phenotype more likely arises predominantly from somatic or extrasynaptic channels, where mechanically activated currents can exert a strong influence on membrane potential and intracellular Ca^2+^ signalling.

### Physiological relevance of mechanical signalling in cortical neurons

Mechanical signalling is increasingly recognised as a general feature of nervous-system physiology rather than a property restricted to peripheral touch receptors. Developing and adult neural cells experience forces generated by tissue growth, cell migration, axon extension, vascular pulsation, cerebrospinal-fluid movement, osmotic volume changes and cell-generated traction. Within this mechanical environment, Piezo1-dependent stiffness sensing contributes to neural-cell migration and fate specification, and to subsequent neuronal differentiation and maturation (Pathak et al., 2014; J. Li et al., 2022; Nourse et al., 2022; Zheng et al., 2023). Age-related stiffening of the central nervous system (CNS) niche suppresses oligodendrocyte progenitor function (Segel et al., 2019), and Piezo1-dependent tissue mechanics in the developing brain can regulate cell adhesion and the long-range distribution of axon-guidance cues, linking local mechanical state to neural patterning (Pillai et al., 2026). Piezo1 therefore operates within a broader mechanochemical framework in which forces regulate not only rapid electrical activity but also developmental, synaptic and transcriptional programmes.

Cortical neurons are particularly relevant to this framework because mechanically gated cation currents recorded at the soma can be large and rapid enough to trigger action potentials in neocortical and hippocampal pyramidal neurons (Nikolaev et al., 2015). Piezo1 has also been implicated in the response of primary cortical neurons to low-intensity ultrasound, where channel inhibition or knockdown reduces Ca^2+^ entry and downstream CaMKII-CREB-c-Fos signalling (Qiu et al., 2019). Our data provide a post-translational mechanism capable of tuning this neuronal response. A lower Piezo1 activation threshold could increase the probability that physiological tissue deformation, local swelling, changes in ECM tension or therapeutic mechanical stimulation produce depolarisation and Ca^2+^-dependent signalling.

Such modulation need not be harmful. Controlled Piezo1 activation can engage neuronal Ca^2+^-dependent signalling, as shown during low-intensity ultrasound stimulation, while astrocytic Piezo1-mediated mechanotransduction supports adult neurogenesis, hippocampal long-term potentiation, learning and memory (Qiu et al., 2019; Chi et al., 2022). Conversely, excessive or sustained Piezo1 activation has been associated with Ca^2+^-calpain-dependent neuronal injury after oxygen-glucose deprivation/reoxygenation and to demyelination and neuronal damage in CNS models (Wang et al., 2019; Velasco-Estevez et al., 2020). The consequences of increased Piezo1 mechanosensitivity are therefore likely depend on the magnitude, duration and spatial distribution of the mechanical stimulus, as well as on the resting excitability and Ca^2+^-buffering capacity of the cell. Thus, a modest increase in mechanosensitivity may facilitate adaptive signalling, whereas sustained or repetitive activation could promote Ca^2+^ overload, aberrant firing or maladaptive gene expression. That hypoglycosylation exerts its largest effect at submaximal stimulation is therefore physiologically important, since it predicts that mechanical inputs normally too weak to elicit a substantial neuronal response could become capable of triggering depolarisation and Ca^2+^-dependent signalling. During neurodevelopment, such a shift in responsiveness could cause normally weak mechanical cues to evoke disproportionately large signals, potentially perturbing neuronal differentiation, circuit wiring and synapse maturation. This mechanism could therefore contribute to the congenital brain abnormalities reported in patients with congenital disorders of glycosylation (CDG) (Paprocka et al., 2021).

### Pathological mechanical stimulation and cortical vulnerability

The same mechanism may become detrimental when the brain is exposed to trauma, oedema, inflammation or marked changes in tissue stiffness. Rapid stretch of cultured cortical neurons produces strain- and strain-rate-dependent membrane permeabilisation (Geddes et al., 2003), and mechanical injury dynamically alters cortical network firing patterns over time (Sullivan et al., 2024). Mechanically activated channels are unlikely to be the only route for injury-induced ion entry, because severe deformation can disrupt the membrane directly and activate multiple ion channels, receptors and intracellular stores. Nevertheless, a Piezo1 population with a reduced activation threshold would be expected to respond earlier during deformation and could amplify the initial Na^+^ and Ca^2+^ load before overt membrane failure occurs.

Several downstream processes could convert this early signal into network dysfunction. Piezo1-mediated inward current can depolarise the neuronal membrane, recruit voltage-gated Na^+^ and Ca^2+^ channels, enhance action potential triggering and neurotransmitter release (Nikolaev et al., 2015), and engage Ca^2+^-dependent kinases and transcription factors (Qiu et al., 2019). In cortical networks, these effects could facilitate synchronised firing or lower the threshold for propagating depolarisation. Piezo1 activation in astrocytes (Chi et al., 2022; Csemer et al., 2024), microglia (Jäntti et al., 2022; Malko et al., 2023), vascular endothelial cells (Wang et al., 2016) and oligodendroglial lineage cells (Velasco-Estevez et al., 2020) can additionally alter gliotransmission, fagocytic and inflammatory responses, vascular tone and myelination. The present experiments were designed around cortical neurons and cannot resolve these multicellular contributions, but because hypoglycosylation would act globally, the *in vivo* phenotype may reflect a convergence of neuronal hyper-responsiveness with altered glial and vascular mechanotransduction.

The substrate dependence observed here is also relevant to disease, because CNS injury and disease remodel the extracellular mechanical environment in ways that could interact with glycosylation state. Glial scar formation is accompanied by marked local tissue softening and changes in laminin and collagen IV (Moeendarbary et al., 2017), and neuroinflammation produces reversible cortical matrix softening that correlates with disease activity (Silva et al., 2024). The mechanical environment also changes with ageing, which increases the stiffness of CNS progenitor niches, and during neurodegeneration, where reduced brain stiffness has been reported in Alzheimer’s disease (Murphy et al., 2011; Segel et al., 2019). Since ECM composition modulates Piezo1 responsiveness to mechanical force (Lai et al., 2022), such disease-associated changes could either enhance or mask glycosylation-dependent differences in channel mechanosensitivity. Glycosylation state and tissue mechanics should therefore not be treated as independent risk factors, since they may interact in a context-dependent manner to determine where, when and at what stimulus intensity mechanosensitive signalling becomes pathological.

### Implications for PMM2-CDG and mechanically triggered neurological episodes

The findings offer a plausible link between defective glycosylation and mechanically triggered neurological deterioration in PMM2-CDG. Stroke-like episodes in this disorder can follow minor head trauma or intercurrent illness and often occur without a vascular lesion that fully explains the neurological deficit (Izquierdo-Serra et al., 2018; Wicker et al., 2023). The substrate dependence of the Piezo1 phenotype may be particularly relevant in this context. The cortical parenchyma that transmits acute traumatic deformations contains a predominantly non-fibrillar extracellular matrix rich in hyaluronan and lecticans, whereas fibrillar collagen is comparatively scarce and is concentrated mainly in the meninges and cerebrovascular basement membranes (Ruoslahti, 1996; Yamaguchi, 2000; Hubert et al., 2009). Thus, although PLL does not reproduce the composition of brain extracellular matrix, the sensitisation observed under non-collagen-mediated adhesion conditions raises the possibility that hypoglycosylated Piezo1 is especially consequential in mechanically challenged parenchymal neurons. By lowering the channel activation threshold, hypoglycosylation could allow mechanical loads that are normally subthreshold to evoke greater cation and Ca^2+^ entry, neuronal depolarisation and network hyperexcitability after otherwise minor cranial trauma.

This hypothesis complements the previously proposed Ca_V_2.1 channelopathy in PMM2-CDG, in which hypoglycosylation of Ca_V_2.1 subunits produces gain-of-function gating changes resembling pathogenic *CACNA1A* variants and has been proposed to contribute to the clinical overlap between PMM2-CDG and *CACNA1A*-associated ataxia and stroke-like episodes (Izquierdo-Serra et al., 2018; Martínez-Monseny et al., 2019). Piezo1 and Ca_V_2.1 could act sequentially. Mechanically evoked Piezo1 current would depolarise cortical neurons, while gain-of-function Ca_V_2.1 channels would enhance voltage-dependent Ca^2+^ entry presynaptically to promote neurotransmitter release. In this model, global hypoglycosylation creates a multichannel disorder in which both the initiating mechanical sensor and the downstream excitability machinery are both biased towards activation, potentially lowering the threshold for network hyperexcitability or spreading-depolarisation-like events after trauma, tissue swelling or inflammation.

Pharmacological inhibition of N-glycan processing provides a useful model of hypoglycosylation but does not fully reproduce PMM2-CDG. PMM2 deficiency affects an earlier step in glycan biosynthesis and can influence both N-glycan occupancy and subsequent maturation across many proteins, whereas swainsonine and kifunensine selectively inhibit downstream glycan-processing steps. The convergence of site-directed mutagenesis and pharmacological inhibition nonetheless demonstrates that Piezo1 function is sensitive to its glycosylation state. Establishing whether this mechanism contributes to PMM2-CDG pathophysiology will require further research using disease-relevant models.

## Conclusions

This study identifies mature N-glycosylation as a previously unrecognised regulator of Piezo1 force sensing. Disrupting N-glycosylation at either conserved cap-domain site, or pharmacological restricting of N-glycan maturation, lowers the mechanical activation threshold without increasing maximal current or slowing inactivation. This gain-of-function phenotype is strongly shaped by the extracellular adhesive environment and, in primary murine cortical neurons, is accompanied by increased somatic Piezo1 abundance and enhanced Ca^2+^ responses to submaximal mechanical stimulation. These findings extend the role of ion-channel glycosylation beyond biosynthesis and trafficking to an active determinant of mechanogating and suggest that glycosylation defects may increase cortical vulnerability by enabling normally weak or innocuous mechanical inputs to more effectively recruit Piezo1-dependent electrical and Ca^2+^ signalling. Together, the results provide a framework for investigating Piezo1 as one component of the altered neuronal mechanotransduction that may accompany PMM2-CDG and other neurological disorders of glycosylation.

## Additional information

### Data availability statement

All data underlying the results and needed to evaluate the conclusions are present in the paper. The original data used and analysed to generate these results are available from the corresponding author upon reasonable request.

### Author contributions

A.E-P. and J.M.F-F. were responsible for the study hypothesis and design of the work. Material preparation, data acquisition and analysis were performed by A.E-P., G.R-U., A.F-A. and J.C-G. A.E-P. and J.M.F-F. drafted the manuscript. All authors commented on previous versions of the manuscript. All authors read and approved the final version of the manuscript submitted for publication.

## Acknowledgements

We thank Ms. C. Plata for excellent technical assistance.

